# Ultrastructural and functional analysis of extra-axonemal structures in trichomonads

**DOI:** 10.1101/2021.07.26.453887

**Authors:** Veronica M. Coceres, Lucrecia S. Iriarte, Abigail Miranda-Magalhães, Thiago André Santos de Andrade, Natalia de Miguel, Antonio Pereira-Neves

**Author notes:** corresponding authors (APN). Fiocruz PE, Instituto Aggeu Magalhães, Departamento de Microbiologia, Av. Professora Moraes rego, s/n - Cidade Universitária, Recife, PE, Brazil. Zip code: 50740-465., (VMC). INTECH, Intendente Marino km 8.2, Chascomús, Provincia de Buenos Aires (CP 7130), Argentina.

## Abstract

*Trichomonas vaginalis* and *Tritrichomonas foetus* are extracellular flagellated parasites that inhabit humans and other mammals, respectively. In addition to motility, flagella act in a variety of biological processes in different cell types; and extra-axonemal structures (EASs) has been described as fibrillar structures that provide mechanical support and act as metabolic, homeostatic and sensory platforms in many organisms. Here, we identified the presence of EASs forming prominent flagellar swellings in *T. vaginalis* and *T. foetus* and we observed that their formation was associated with the parasites adhesion on the host cells, fibronectin, and precationized surfaces; and parasite:parasite interaction. A high number of rosettes, clusters of intramembrane particles that has been proposed as sensorial structures, and microvesicles protruding from the membrane were observed in the EASs. The protein VPS32, a member of the ESCRT-III complex crucial for diverse membrane remodeling events, the pinching off and release of microvesicles, was found in the surface as well as in microvesicles protruding from EASs. Moreover, we demonstrated that overexpression of VPS32 protein induce EAS formation and increase parasite motility in semi-solid medium. These results provide valuable data about the role of the flagellar EASs in the cell-to-cell communication and pathogenesis of these extracellular parasites.

## INTRODUCTION

The eukaryotic flagella are highly conserved microtubule-based organelles that extend from the cell surface. These structures, beyond being essential for cell locomotion and movement of fluids across the tissues and cells, are signaling platforms that receive and send information to drive cellular responses (Akella et al., 2020; Carter & Blacque, 2019). These functions are crucial for health, development and reproduction processes in most eukaryotes, including humans (Anvarian et al., 2019; Wan & Jekely, 2020). In addition to cell movement (Imhof et al., 2019) and sensory functions (Maric et al., 2010), a variety of microorganisms employ flagella to control feeding (Dolger et al., 2017), mating (Fussy et al., 2017), cytokinesis (Hardin et al., 2017; Ralston et al., 2006), cell morphogenesis (Vaughan, 2010), cell communication (Szempruch et al., 2016) and cell adhesion (Frolov et al., 2018). Among these microorganisms, there are important human and veterinary parasitic protists, i.e. trichomonads, trypanosomatids, diplomonads and apicomplexa that exert a devastating economic burden on global healthcare systems and agriculture (Kruger & Engstler, 2015).

The trichomonads (Metamonada, Parabasalia) *Trichomonas vaginalis* and *Tritrichomonas foetus* are extracellular parasites that inhabit humans and other mammals, respectively. *T. vaginalis* is responsible for trichomoniasis, the most common non-viral sexually transmitted infection in men and women (WHO, 2018). Most infected people are asymptomatic, but when symptoms do occur, they can range from mild irritation to severe inflammation in various regions of the reproductive tract (Van Gerwen & Muzny, 2019). *T. vaginalis* is also associated with pelvic inflammatory disease, pregnancy complications, preterm birth, infertility (Kissinger, 2015; Meites et al., 2015) as well as increased risk to HIV (McClelland et al., 2007; Van Der Pol et al., 2008), papillomavirus infection and cervical or prostate cancer (Gander et al., 2009; Stark et al., 2009; Sutcliffe et al., 2009; Twu et al., 2014). *T. foetus* is a widespread pathogen that colonizes the reproductive tract of cattle and the large intestine of cats, leading to bovine and feline tritrichomonosis, respectively. Bovine tritrichomonosis is a venereal infection that causes significant economic losses in beef and dairy farming due to early embryonic death, abortion and infertility or culling of parasite carriers (Mardones et al., 2008; Martin-Gomez et al., 1998). Feline tritrichomonosis causes chronic diarrhea in cats (Gookin et al., 2017). *T. foetus* also lives as a commensal in the nasal and gastrointestinal mucosa of pigs (Dabrowska et al., 2020).

In each trichomonads genus, the flagella vary in number and size: *T. vaginalis* and *T. foetus* have five and four flagella, respectively (Benchimol, 2004). Like most eukaryotes, the structural basis of the trichomonads motile flagella is the canonical ‘9+2’ microtubular axoneme surrounded by plasma membrane (Benchimol, 2004). In both species, the plasma membrane of the anterior flagella has rosette-like formations that have been proposed as sensorial structures (Benchimol et al., 1982; Honigberg et al., 1984). Based on this, some authors have suggested that the flagella could be involved in migration and sensory reception in trichomonads during adherence to host tissue and amoeboid morphogenesis (de Miguel et al., 2012; Kusdian et al., 2013; Lenaghan et al., 2014). However, the flagellar role during parasite cell adhesion, amoeboid transformation and cell-to-cell communication is still poorly understood.

In other organisms, flagella can send information via ectosomes (also called microvesicles), a type of extracellular vesicle that protrude and shed from the cell surface (Wang & Barr, 2018). In *Trypanosome*, these ectosomes can transfer virulence factors from one parasite to the other contributing to the pathogenesis (Szempruch et al., 2016). In this sense, our group recently reported that *T. vaginalis* release flagellar ectosomes that might have an important role in cell communication (Nievas et al., 2018). Proteins from the endosomal sorting complex required for transport (ESCRT) machinery are involved in flagellar ectosomes release in protists. Specifically, ESCRT-III proteins may play a central role in promoting ectosome budding from the flagellum membrane (Long et al., 2016). However, the localization and possible functions of ESCRT proteins in the trichomonads flagella have not been determined yet.

In addition to axoneme and ectosomes, the assembly of extra-axonemal structures (EASs) occurs in many organisms ranging from mammalian and insects (Miao et al., 2019; Zhao et al., 2018) to protists, e.g. euglenozoa, dinoflagellates and *Giardia* (Maia-Brigagao et al., 2013; Moran, 2014; Portman & Gull, 2010). EASs are evolutionarily convergent, highly organized fibrillar structures that provide mechanical support and act as metabolic, homeostatic and sensory platforms for the regulation of flagellar beating (Moran, 2014; Portman & Gull, 2010). Depending on the cell type, EASs can be symmetrically or asymmetrically arranged around the axoneme and they can run along almost the entire length or only a portion of the flagellum (Portman & Gull, 2010). In protists, the paraflagellar rod (PFR), which is seen in trypanosomatids, is the best characterized EAS. PFR is required for motility, parasite attachment to host cells, morphogenesis and cell division (Portman & Gull, 2010). Although EASs, formed by thin filaments, have been described in some trichomonads and related parabasalid species (G. Brugerolle, 1999, 2005; Brugerolle & König, 1994; Mattern et al., 1973), there are no reports on the existence and role of EASs in *T. vaginalis* and *T. foetus.* In this work, using a detailed ultrastructural analysis, we identified the presence of EASs forming prominent flagellar swellings in *T. vaginalis* and *T. foetus*. Interestingly, we found that the formation of flagellar swellings was associated to: (a) amoeboid morphogenesis; (b) adhesion to host cells and fibronectin; and (c) parasite: parasite interaction. A high number of rosettes and microvesicles protruding from the membrane can be found in the EAS. Finally, we found that overexpression of a member of the ESCRT-III complex that localized at the flagellar swelling, named VPS32, induce EAS formation and increase parasite motility in semi-solid medium. Our data highlight a role for the EAS in the cell-to-cell communication and pathogenesis in *T. vaginalis* and *T. foetus*.

## RESULTS

### Presence of flagellar swellings in *T. vaginalis* and *T. foetus*

To examine in detail the trichomonads flagellar morphology, we initially observed three wild-type strains of *T. vaginalis* and two different strains of *T. foetus* grown axenically using scanning (SEM) and transmission (TEM) electron microscopy. As can be visualized in Figure 1, *T. vaginalis* has four anterior flagella (AF); *T. foetus* has three AF; both parasites have one recurrent flagellum (RF) that forms the undulating membrane. In *T. vaginalis,* the RF runs along two-thirds of the cell and no free portion is developed, whereas in *T. foetus* the RF reaches the posterior end of the cell and extends beyond the undulating membrane as a free tip (Fig. 1). As expected, the flagella of most of the parasites (between 89 to 99%) displayed a classical ultrastructure: a diameter of 250-300 nm along their length and the flagellar membrane around the “9+2” axoneme (Fig. 1, insets). However, the presence of flagellar swellings in the tip or along the AF and RF was observed in 1-11% of *T. vaginalis* and 2-5% of *T. foetus* parasites analysed by SEM (Fig 2A). These swellings exhibited two different morphologies: “sausage-like” and “spoon-like” (Fig. 2B). The “sausage-like” swelling runs laterally or surrounding the axoneme, exhibiting a range size from 0.1 to 1 µm in thickness and a variable-length from 0.3 to 6 µm in *T. vaginalis* and up to 1 µm in *T. foetus* (Fig. 2B; Suppl Fig. 1). In the “spoon-like” swelling, the flagellum wraps around the swelling to form a rounded or ellipsoid structure measuring between 0.5 to 2.5 µm in the major axis in *T. foetus* and more than 4 µm long in *T. vaginalis* (Fig. 2B; Suppl Figs. 2A-C). The “spoon-like” structure can exhibit a flattened or concave surface in a frontal view, and an aligned, curved, or convex appearance in a side view (Suppl Figs. 2D-H). Based on the results in the Fig. 2A, subsequent experiments were performed using the B7RC2 and K strains of *T. vaginalis* and *T. foetus*, respectively. Curiously, while the “sausage-like” structure was more frequently found in *T. vaginalis*, the “spoon-like” was more common in *T. foetus* (Fig. 2C). Interestingly, although these structures can be found in all flagella, they are more frequent in AF in *T. vaginalis* and RF in *T. foetus* (Fig. 2D). The analysis of “spoon-like” and “sausage-like” flagellar distribution demonstrate that both types of structures can be identified in the RF and AF in *T. foetus* as well as in the AF of *T. vaginalis* (Fig. 2E). However, only “sausage-like” structures were detected in the RF of *T. vaginalis* (Fig. 2E-F). In *T. foetus*, around 3-6% of flagella with swelling exhibited “sausage” and “spoon-like” structures in the same flagellum (Fig. 2E-G). When the relative position of both types of structures along the flagella was evaluated, we noted that the “sausage-like” swelling was predominantly found at the flagellar tip of *T. vaginalis* and AF of *T. foetus* (Fig. 2H); however, it was also observed in the middle (Figs. 2H-I) and, rarely, at the tip and in the middle of the same flagellum (Figs. 2H-J). The “spoon-like” structure was usually located at the AF’s tip of both parasites, and, occasionally, seen in the middle of *T. foetus* RF (Figs. 2K-L).

**Figure 1.**
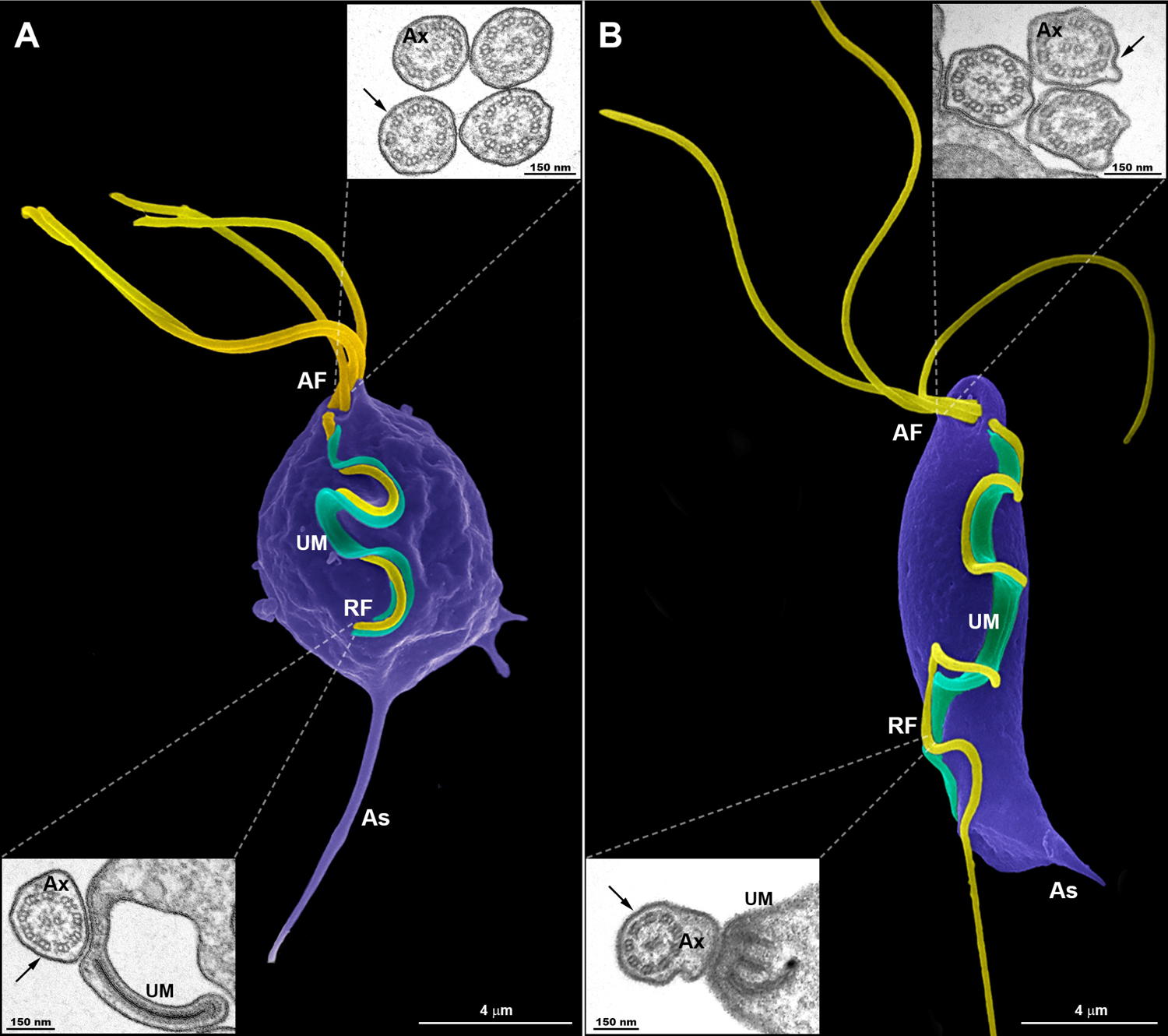
Typical morphology of trichomonads grown in axenic culture. SEM of *T. vaginalis* (**A**) and *T. foetus* (**B**) with the pear-shaped cell bodies colored violet and flagella colored yellow. *T. vaginalis* exhibits four anterior flagella (AF), whereas *T. foetus* has three AF; both parasites have one recurrent flagellum (RF) that runs posteriorly along the cell body, forming an undulating membrane (UM – colored green). The T. *vaginalis*-RF is shorter than *T. foetus*-RF. The later displays a distal free end. The flagella are the same width along their length and no swellings or enlarged areas are seen. The axostyle (As) tip is visible. The insets are TEM images of the AF (upper insets) and RF (lower insets) in representative transverse sections, viewed from the proximal and distal end, respectively. Note the 9+2 axoneme (Ax) enclosed within the flagellar membrane (arrows). No EAS are seen.

**Figure 2.**
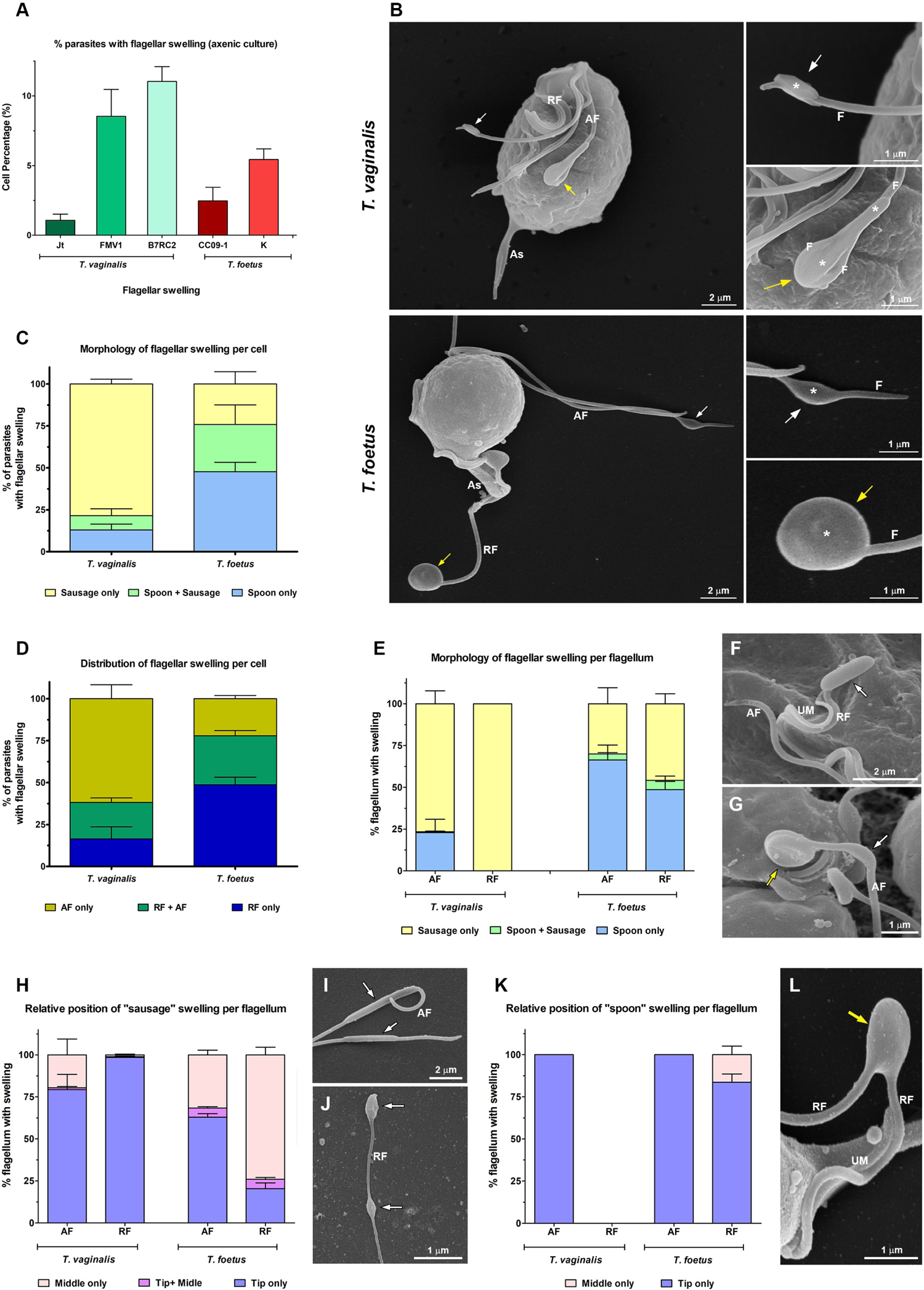
Morphological analyses of flagellar swellings in *T. vaginalis* and *T. foetus* under standard growth conditions. (**A**) Quantification of the percentage of parasites that display flagellar swellings. The values are expressed as the means ± the standard deviation (SD) of three independent experiments, each performed in duplicate. 500 parasites per sample were randomly counted. (**B**) General and detailed views of flagellar swellings (*) in *T vaginalis* and *T. foetus* obtained by SEM. The swellings can exhibit two different morphologies: “sausage shaped” (white arrows) and “spoon shaped” (yellow arrows). Notice that the “sausage-like” swelling runs laterally to the flagellum (F), whereas, in the spoon shaped structure, the swelling is surrounded by flagellum. AF, anterior flagella; RF, recurrent flagellum; As, axostyle. (**C-D**) Quantitative analysis of morphology (**C**) and distribution (**D**) of flagellar swellings per parasite. Three independent experiments in duplicate were performed and 100 parasites exhibiting at least one swelling were randomly counted per sample using SEM. Data are expressed as percentage of parasites with flagellar swelling ± SD. AF, anterior flagella; RF, recurrent flagellum. (**E**) Quantification of the morphology of flagellar swelling per flagellum. The values are expressed as the means of the percentage of flagellum with swelling ± SD of three independent experiments, each performed in duplicate. 100 anterior and recurrent flagella with swelling per sample were randomly counted using SEM. AF, anterior flagella; RF, recurrent flagellum. (**F-G**) Detailed views of a recurrent flagellum (RF) of *T. vaginalis* (**F**) and an anterior flagellum (AF) of *T. foetus* (**G**) by SEM. UM, undulating membrane. In (**F**), a sausage shaped swelling (arrow) is seen in the tip of the flagellum. Notice in (**G**) the presence of “sausage” (white arrow) and “spoon-like” (yellow arrow) structures in the same flagellum. (**H**) Analysis of the relative position of “sausage” swelling per flagellum. Three independent experiments in duplicate were performed and 100 anterior and recurrent flagella with swelling per sample were randomly counted using SEM. Data are expressed as percentage of flagellum exhibiting swelling ± SD. AF, anterior flagella; RF, recurrent flagellum. (**I-J**) SEM of sausage shaped structures (arrows) located along the anterior flagella (AF) of *T. vaginalis* (**I**) and at the tip and in the middle of the same recurrent flagellum of *T. foetus* (**J**). (**K**) Quantification of the relative position of “spoon” swelling per flagellum. The values are expressed as the means of the percentage of flagellum exhibiting swelling ± SD of three independent experiments, each performed in duplicate. 100 anterior and recurrent flagella with swelling per sample were randomly counted using SEM. AF, anterior flagella; RF, recurrent flagellum. (**L**) SEM of a spoon shaped structure (arrow) located in the middle of *T. foetus* recurrent flagellum (RF). UM, undulating membrane.

### Flagellar swellings are extra-axonemal structures (EASs) formed by thin filaments

To investigate the ultrastructural characteristics of flagellar swellings in trichomonads, we analyzed the flagella using negative staining and ultrathin section techniques for transmission electron microscopy (TEM) (Figs. 3-4). Our results demonstrate that flagellar microtubules are surrounded by a continuous membrane that comes from the cell body and that “sausage-like” swelling is formed by thin extra-axonemal filaments that run longitudinally along the axonemes (Fig. 3). A detailed analysis of longitudinal and transverse sections showed that the extra-axonemal filaments measure around 3 - 5 nm in diameter and their length varies according to the length of the swelling (Fig. 3C). To further understand the morphological organization of “sausage” swelling, we analysed complementary images acquired in different perspectives (Suppl. Fig. 3). Those results confirmed that the extra-axonemal filaments partially surround the axoneme, although SEM top view images may lead to misinterpretation of the flagella is totally surrounded by the swelling (Suppl. Fig. 3). In an oblique view, we noticed that the axoneme is in a slit of the swelling as a hot dog shaped structure (Suppl. Fig. 3). The “sausage” structures located in the middle of flagella and in the recurrent flagellum are also formed by extra-axonemal filaments (Suppl. Figs. 4-5), indicating that the flagellar swellings are EASs.

**Figure 3.**
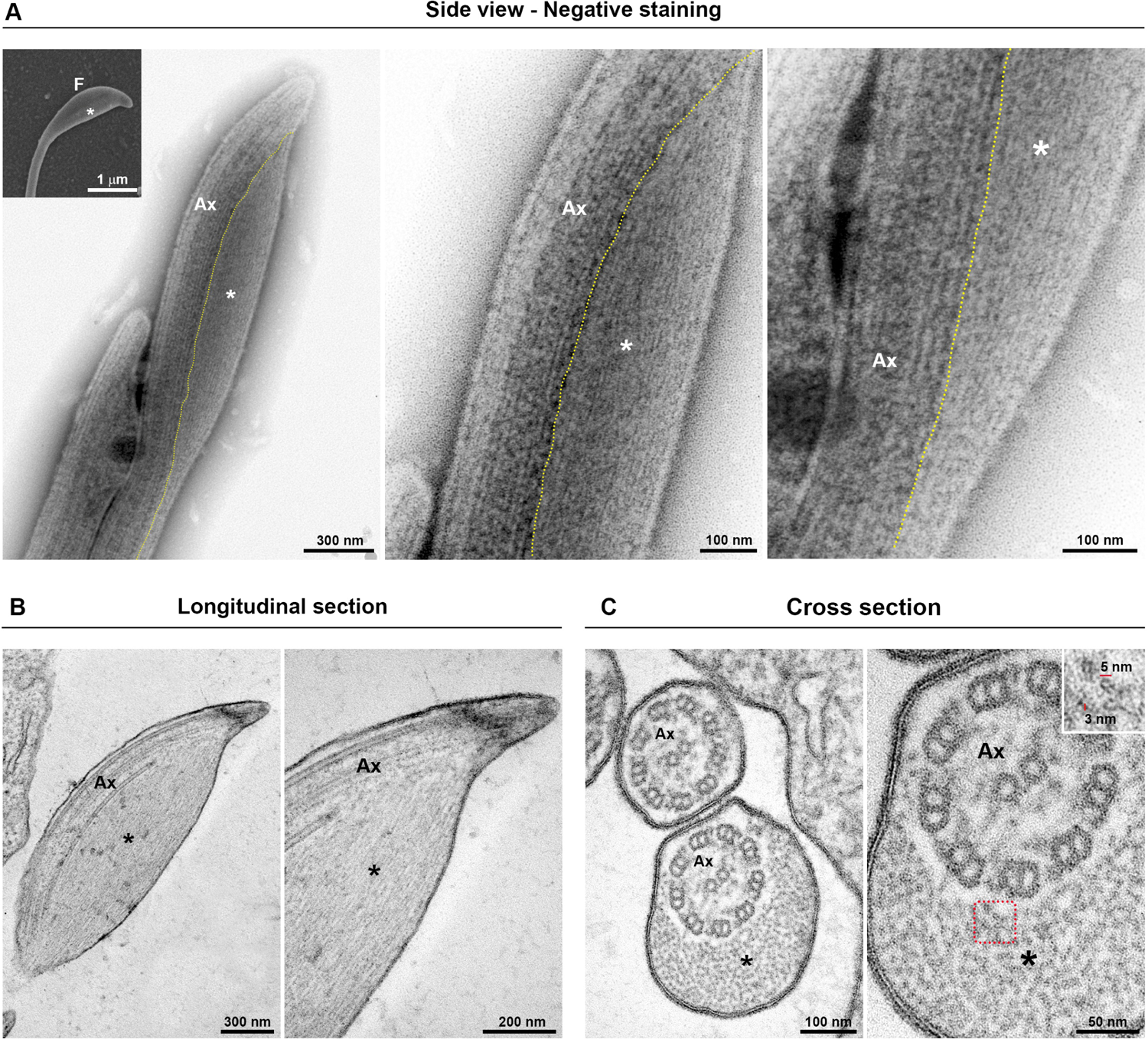
Ultrastructure of the flagellar sausage shaped swelling. The structure is formed by thin extra-axonemal filaments (*) that run longitudinally along the axoneme (Ax). (**A**) Negative staining images of a swelling on side view. The dotted lines indicate boundary between axoneme and the extra-axonemal filaments. Inset, a complementary SEM image is used as reference. F, flagellum. (**B-C**) Longitudinal and cross ultrathin sections. The extra-axonemal filaments measure around 3 - 5 nm in diameter (inset).

**Figure 4.**
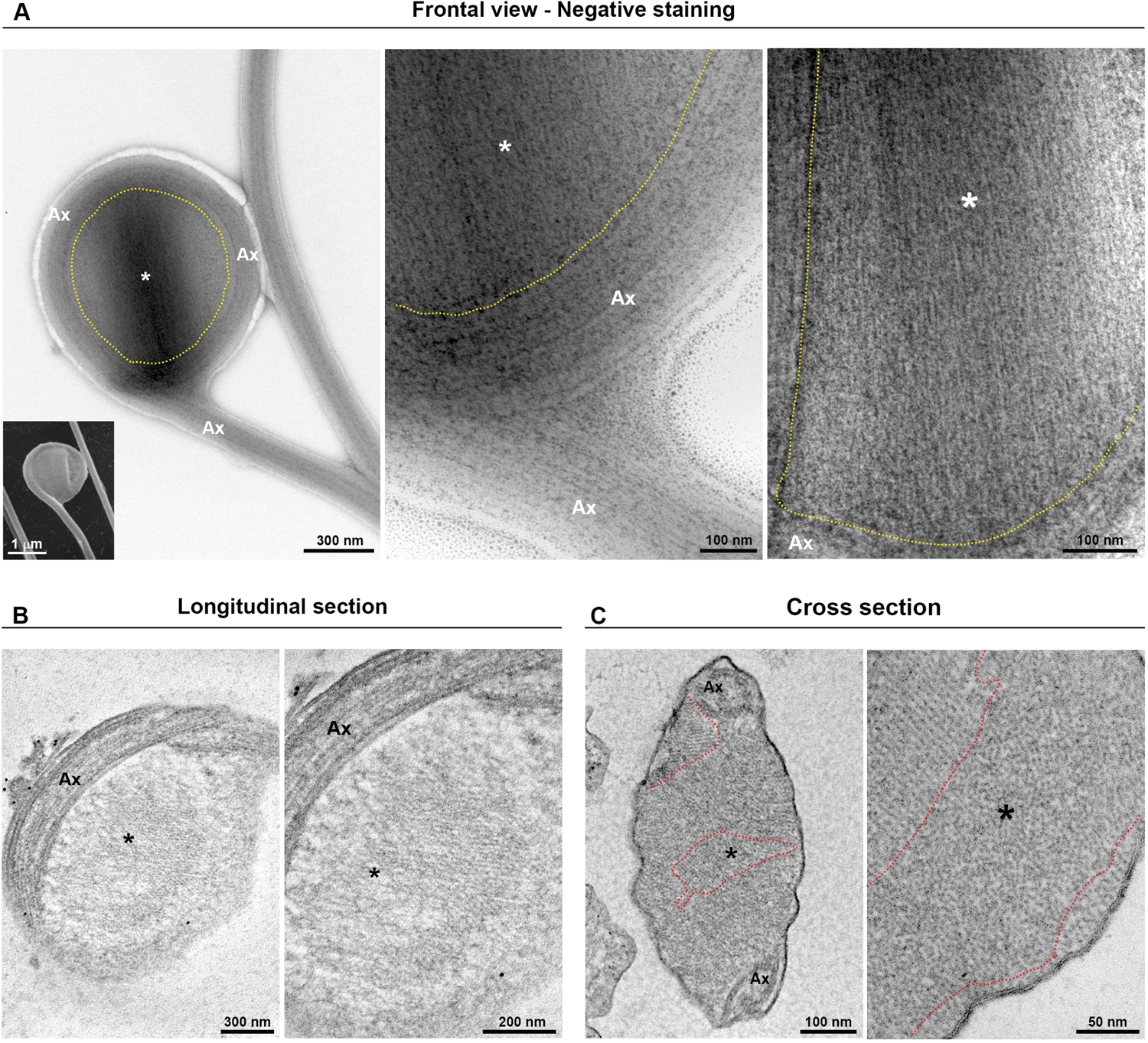
Fine structure of the flagellar “spoon-like” swelling. The structure is formed by folding the axoneme (Ax) around the thin extra-axonemal filaments (*). (**A**) Negative staining images of a swelling on frontal view. The dotted lines indicate boundary between axoneme and the extra-axonemal filaments. Inset, a complementary SEM image is used as reference. (**B**) Longitudinal ultrathin sections. The extra-axonemal filaments display a lattice-like arrange. (**C**) Cross ultrathin sections. The filaments are seen organized in different orientations, as indicated by the dotted lines.

**Figure 5.**
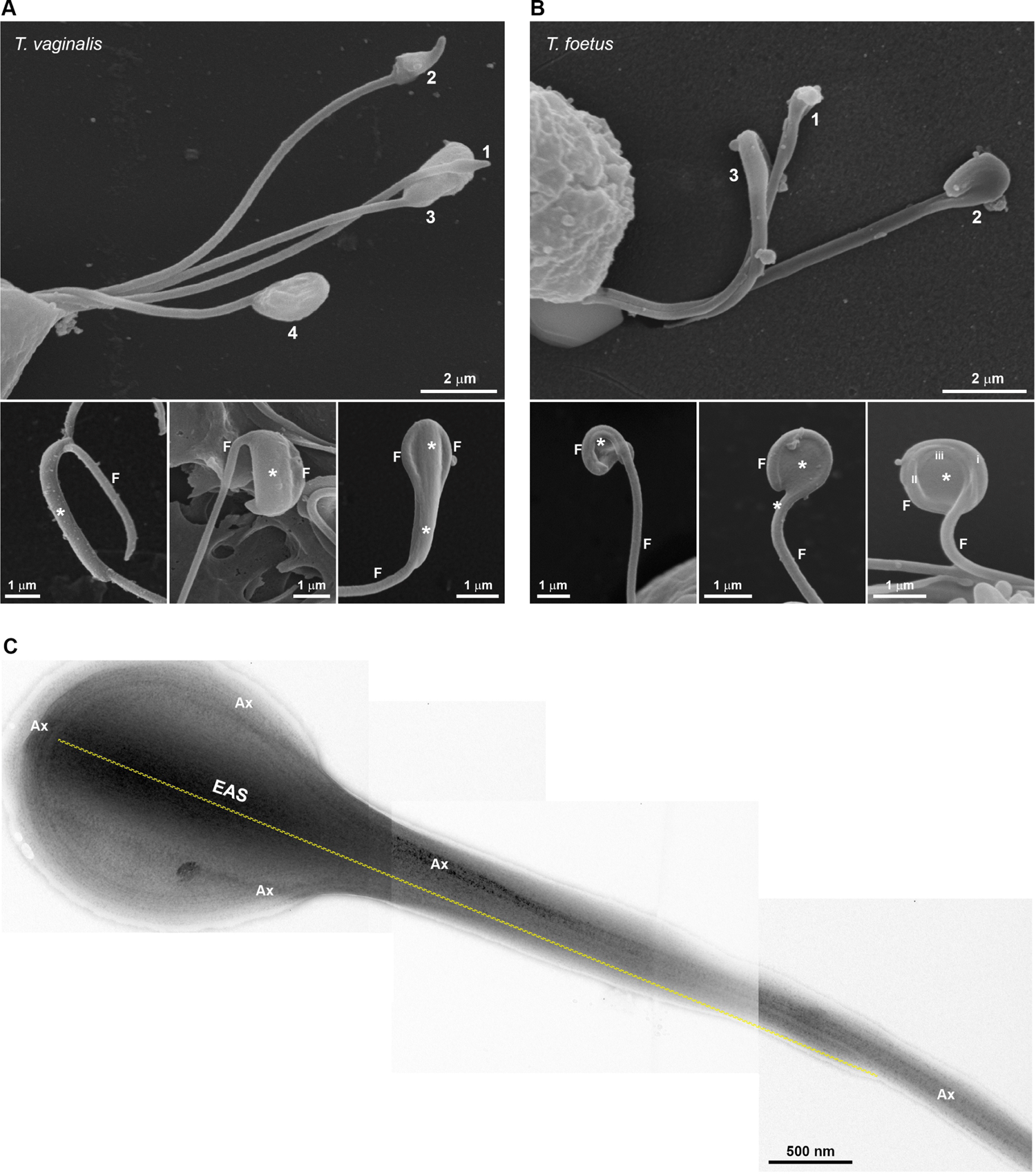
Morphological diversity of flagellar swelling. (**A-B**) SEM showing swellings (*) of different sizes in *T. vaginalis* (**A**) and *T. foetus* (**B**). The numbers 1, 2, 3 and 4 and images suggest plausible stages for the spoon shaped structure formation. In (**B**), the roman numbers (i, ii and iii) indicate the amount of flagellum (F) folds around the swelling. (**C**) Negative staining of a sausage shaped extra-axonemal structure (EAS) surrounding by axoneme (Ax).

Additionally, we demonstrated that the “spoon-type” swelling is also an EASs formed by folding the axoneme around the extra-axonemal filaments (Fig. 4A). When observed in longitudinal sections, the filaments display a lattice-like arrange (Figs. 4B). In a transversal view, it can be observed that the filaments are organized in different orientations (Figs. 4C); probably due to the turns of the axoneme around the filaments. As swellings are formed by extra-axonemal filaments, it is very likely that morphological differences could be attributed to different phases of a single process. Supporting this, SEM analysis suggests that the “sausage” and the “spoon” could be different stages of a single event (Figs. 5A-B). The process might start with a small sausage-shaped EAS that give raise to a “spoon” when the flagella fold around an enlarged EAS and on themselves (Figs. 5A-B). TEM images confirmed that sausage shaped EAS is surrounding by axoneme (Fig. 5C). Because the *T. vaginalis*-RF has no free portion, this could help to explain why only sausage shaped EASs are observed in that flagellum, whereas both sausage and spoon shaped EASs are found in the free tip of *T. foetus*-RF (Suppl. Fig. 6).

The existence of rosette-like formations (clusters of intramembrane particles), proposed as sensorial structures, has been reported in the AF of *T. vaginalis* and *T. foetus* (Benchimol & De Souza, 1990; Benchimol et al., 1981). In this regard, we evaluated the presence of rosettes in the *T. vaginalis* EASs by negative staining technique. Interestingly, we observed that flagella with EASs showed a higher number of rosettes/µm^2^ than those flagella without such structures (Fig. 6). In summary, our results demonstrated that EAS in trichomonads are membrane expansions with different morphologies (sausage/spoon), formed by thin filaments and a high number of rosettes in their membranes.

**Figure 6.**
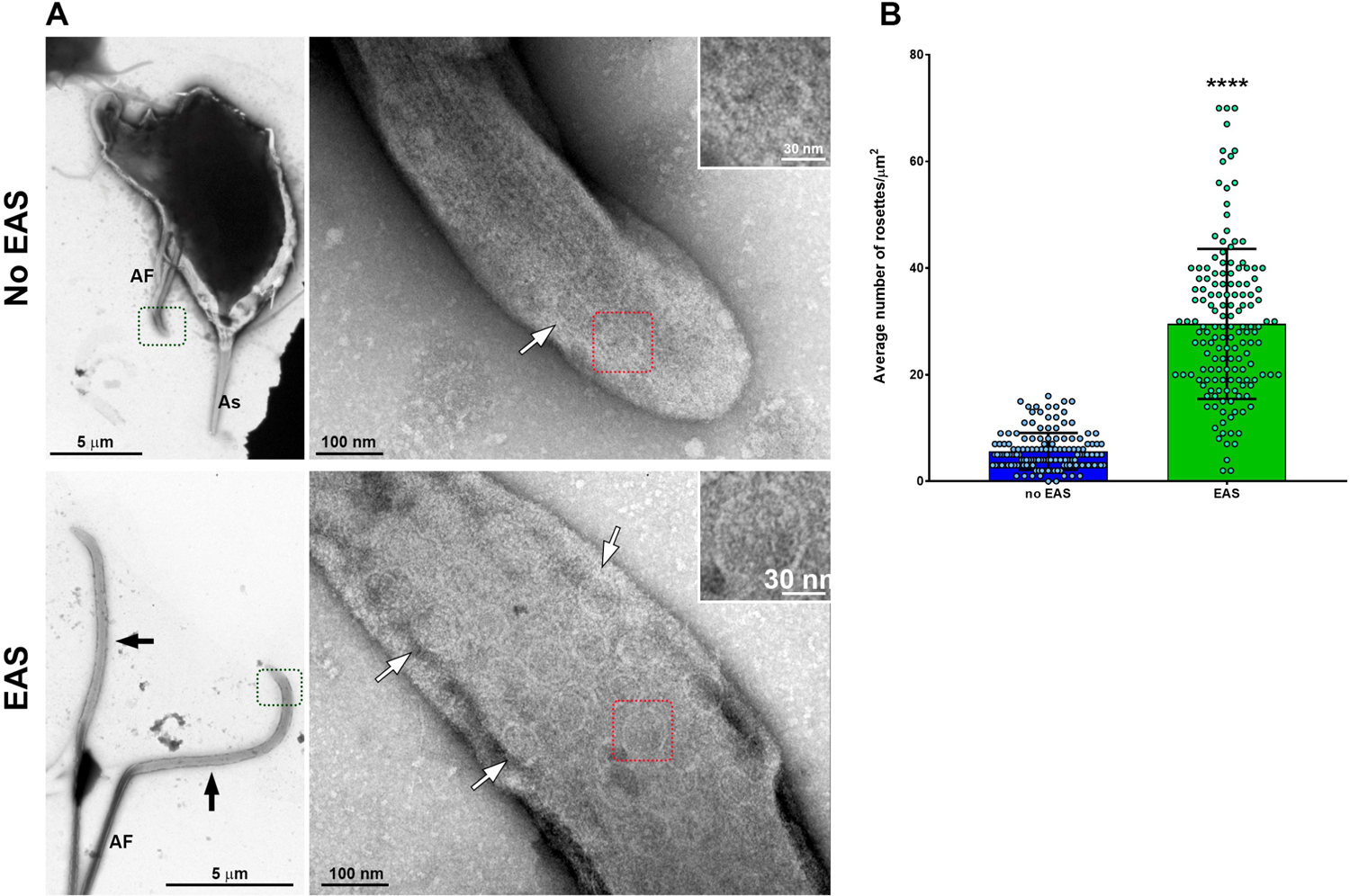
Flagella with swelling exhibit a higher number of rosette-like formations. (**A**) Representative general and detailed views of *T. vaginalis* anterior flagella (AF) without and with extra-axonemal structures (EAS) obtained by TEM. Many rosette-like formations (white arrows) are seen in the flagella with swelling (black arrows). As, axostyle. (**B**) Quantification of the number of rosettes/µm^2^. The columns represent the average number of rosettes/µm^2^ ± the standard deviation (SD) of three independent experiments. 50 flagella with or without swellings per sample were randomly counted using TEM. The dots indicate the values obtained for each flagellum. Flagella with EAS show a higher number of rosettes/µm^2^ than those flagella without EAS. **** p<0.0001 compared to “no-EAS” group using non-parametric t-test (Mann-Whitney test).

### The EASs formation increase during *T. vaginalis* and *T. foetus* attachment process

The ability of trichomonads to colonize the epithelia has been studied in recent years; however, the role of flagella in this process is not fully understood. To evaluate a possible contribution of extra-axonemal structures to parasite attachment, parasites were incubated on fibronectin-coated coverslips or Alcian blue precationized coverslips, washed with PBS to remove non-attached cells, and the formation of EASs was evaluated by SEM (Fig. 7 and Suppl. Fig. 7). Attached parasites remained on the coverslips, whereas non-attached cells were harvested by centrifugation and analysed using SEM. For control, parasites were incubated on uncovered coverslips, collected with a pipette, harvested by centrifugation, and also prepared for SEM. As expected, cells were in suspension and unattached on the uncovered coverslips (not shown); therefore, here, “Control” is defined as non-adherent, suspended cells from uncovered coverslips, whereas non-adherent parasites from fibronectin and Alcian blue interaction assays are called “Non-attached”. Parasites from control exhibit the typical pyriform body and no cell clusters (suppl. Fig. 7A). As expected, attached parasites on fibronectin-coated coverslips exhibited an amoeboid morphology and many flagellar swellings (Fig. 7A). The percentage of fibronectin-Attached parasites with EAS is higher when compared to the Non-attached and control groups (Figs. 7B-C). In control, EASs are found in 9.9% and 3.9% of *T. vaginalis* and *T. foetus*, respectively, whereas EAS formation is observed in 48.9% and 54.6% of fibronectin-Attached *T. vaginalis* and *T. foetus* groups, respectively (Figs. 7B-C). When the parasites were incubated onto coverslips pre-treated with Alcian-blue, the cells were found clustered, mainly *T. vaginalis*, displaying an amoeboid or ellipsoid form in both Attached and Non-attached groups (Suppl. Fig. 7A). Similarly, the percentage of parasites with EAS in the Alcian blue-attached parasite is higher s when compared to control (Suppl. Figs. 7B-C). In control, EASs are found in 12.5% and 5.2% of *T. vaginalis* and *T. foetus*, respectively, whereas EAS formation is observed in 41.7% and 40.2% of Alcian-blue attached *T. vaginalis* and *T. foetus* groups, respectively (Suppl. Figs. 7B-C). Unexpectedly, the percentage of *T. vaginalis* with EAS in the Alcian blue-Non-attached group was significantly higher when compared to control (Suppl. Figs. 7B-C)

**Figure 7.**
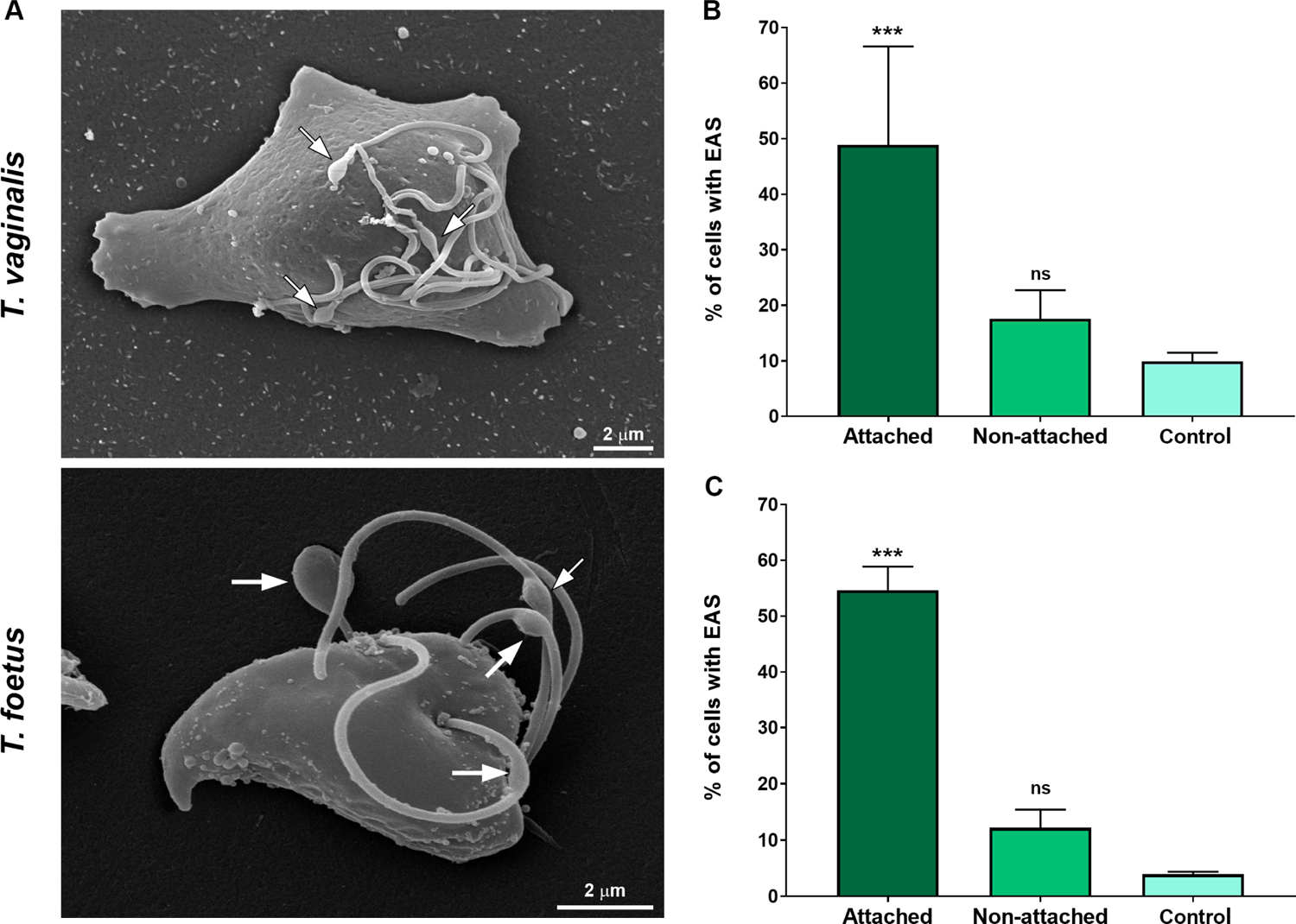
The EASs formation increase during trichomonads attachment on Fibronectin-coated coverslips. (**A**) SEM of *T. vaginalis* and *T. foetus* after adhesion assay on fibronectin-coated coverslips. Arrows indicate the extra-axonemal structures (EAS). Notice that parasites display an amoeboid morphology. (**B-C**) Quantitative analyses in *T. vaginalis* (**B**) and *T. foetus* (**C**). The percentage of cells with EASs was determined by counting of 500 parasites per sample using SEM. Data are expressed as means of three independent experiments in duplicate ± SD. Attached and non-attached: parasites resuspended in PBS incubated on fibronectin-coated coverslips in humidity chamber for 2 h at 37°C and rigorously washed with PBS to remove non-attached cells. Attached parasites remain on the coverslips even after several washes. Non-attached parasites were collected with a pipette, harvested by centrifugation, and prepared for SEM. Control, parasites incubated on uncovered coverslips under the same conditions mentioned above, collected with a pipette, harvested by centrifugation, and prepared for SEM. “Control” is formed by non-adherent, suspended cells from uncovered coverslips, whereas non-adherent parasites from fibronectin are called “Non-attached”. The percentage of parasites displaying EASs is significantly higher in Attached group when compared to Non-attached and control groups. ***p<0.001 compared to control group using One-Way ANOVA test (Kruskal-Wallis test; Dunn’s multiple comparisons test). ns, non-significant.

Next, to evaluate if EASs have a role in host cell interaction, parasites were incubated with host cells and the number of parasites that contain flagellar swellings was quantified using SEM (Fig. 8). Two different ratios of parasites: host cells were used and parasites in absence of host cells were used as control (PBS). Upon exposure, EAS are found in some parasites and some swellings are seen in direct contact with the host cells (Fig. 8A). When *T. vaginalis* parasites are incubated with VECs (vaginal epithelial cells) at 1:1 and 5:1 ratio, the formation of EASs was observed in 27.4% and 30.7% of parasites, respectively (Fig. 8B). Similarly, when *T. foetus* are exposed to PECs (bovine preputial epithelial cells), EASs are observed in 25.6% and 36.6% of attached parasite at ratio of 1:1 and 5:1 respectively (Fig. 8C). Moreover, we observed that these structures were present in flagella of parasites in contact to prostatic cells, preputial mucus content and in contact to bacteria present in the microbiota of the reproductive system (Suppl. Fig. 8). Together, these results indicate that extra-axonemal structures are being formed in response to host cell exposure.

**Figure 8.**
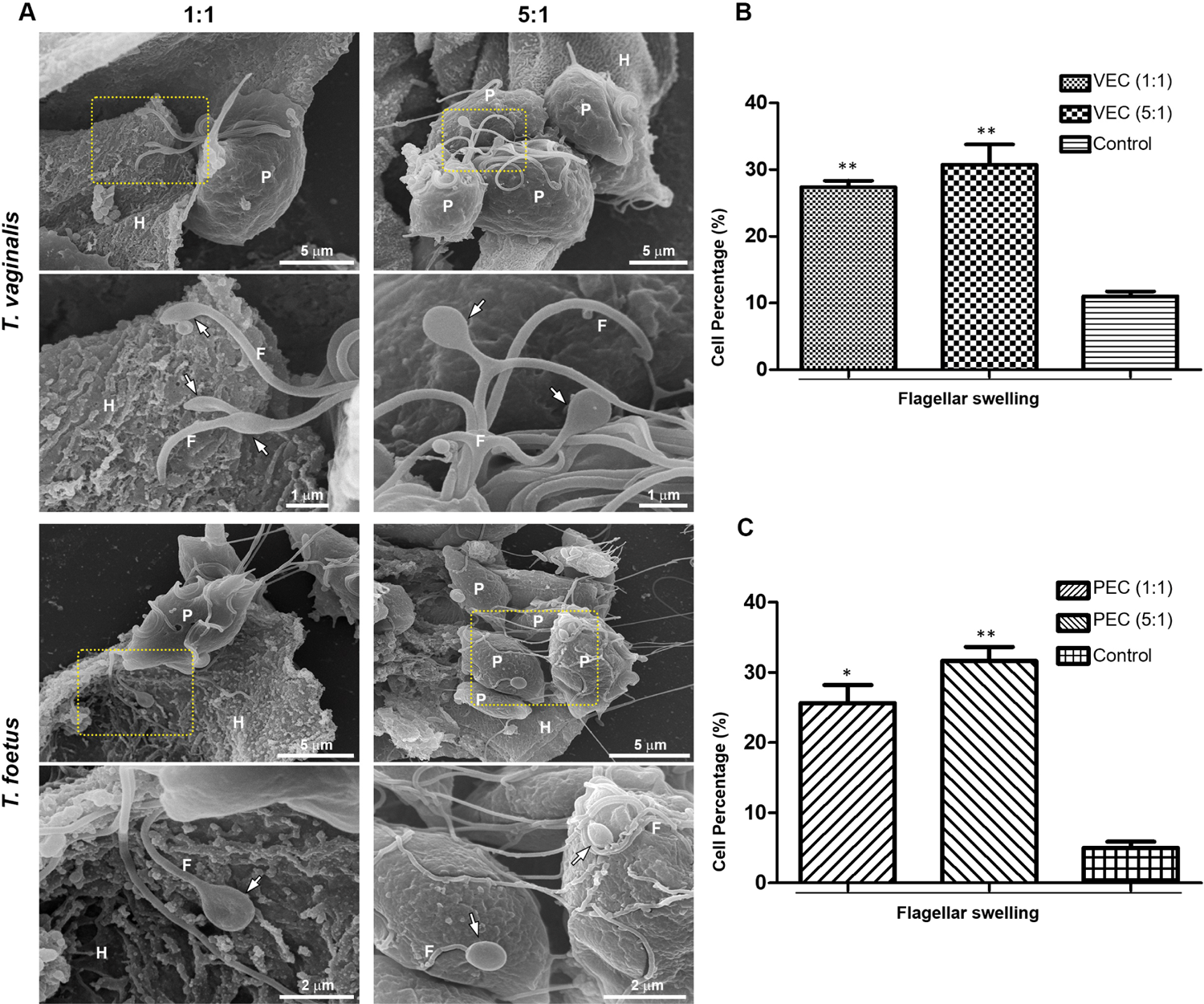
EASs are formed in response to host cell exposure. (**A**) Representative SEM images of *T. vaginalis* and *T. foetus* after host cell interaction. Human vaginal epithelial cell (VECs) and bovine preputial epithelial cell (PECs) were co-incubated with *T. vaginalis* and *T. foetus*, respectively, at cell ratios of 1:1 or 5:1 parasite:host cell in PBS-F (PBS with 1% FBS at pH 6.5) at 37°C for 30 min. Flagellar swelling (arrows) are seen in some parasites (P). Notice the swellings in direct contact to the host cells (H) in the images of 1:1 ratio. (**B-C**) Quantification of the percentage of *T. vaginalis* (**B**) and *T. foetus* (**C**) with flagellar swelling after the host cell interaction. Three independent experiments in duplicate were performed and 500 parasites were randomly counted per sample using SEM. Data are expressed as percentage of parasites ± SD. For the control experiments, parasites incubated in PBS in the absence of host cells were analysed. The percentage of parasites with flagellar swelling increases after the hot cell exposure when compared to control (PBS). * p<0.05; **p<0.01 compared to control using One-Way ANOVA test (Kruskal-Wallis test; Dunn’s multiple comparisons test).

### Microvesicles are shed from the membrane of EASs

Flagella can send information through microvesicles (MVs) released from their membranes (Szempruch et al., 2016; Wood et al., 2013). Previous results from our group demonstrated that *T. vaginalis* release flagellar microvesicles; although their biological relevance still is unknown (Nievas et al., 2018). Here, we observed the presence of MVs associated to EASs by SEM, negative staining and ultrathin sections (Figs. 9A). We demonstrated that 44.1% and 47.1% of *T. vaginalis* and *T. foetus* from axenic culture with flagellar swelling, respectively, exhibit MVs protruding from the flagellar membrane of the EASs (Fig. 9B). Considering that formation of EASs increase during parasite attachment to host cells, the presence of MVs in EASs membrane and the role of MVs in cell communication, these results suggest a possible role of MVs protruding from EASs in attachment or parasite:host cell communication.

**Figure 9.**
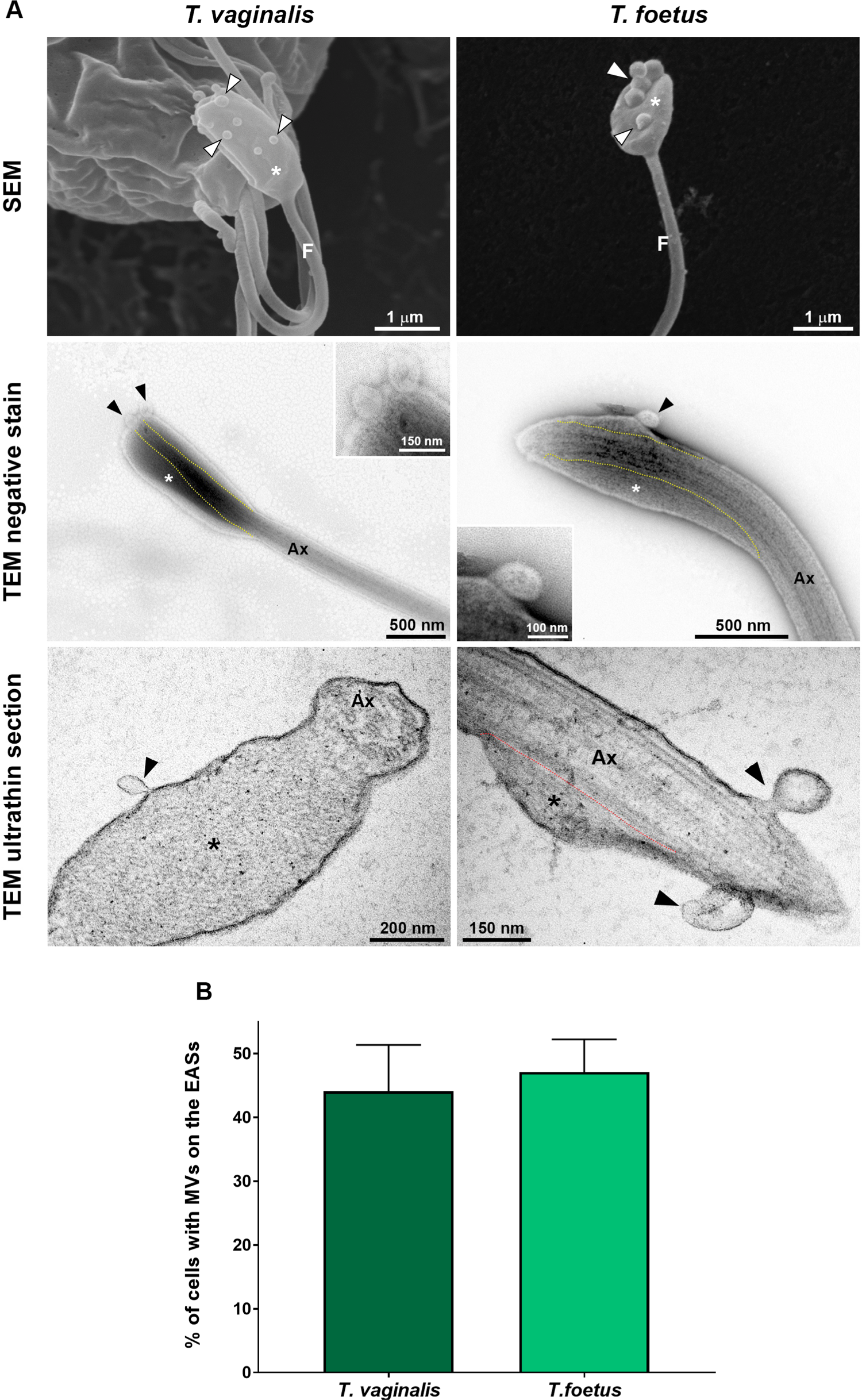
EASs release microvesicles-like structures. (**A**) Representative micrographs of MVs (arrowheads) protruding from the flagellar membrane of the EASs (*) of *T. vaginalis* and *T. foetus*. The images were obtained by SEM (first row), negative staining (second row) and ultrathin sections (third row). The dotted lines indicate boundary between axoneme (Ax) and the extra-axonemal filaments (*). (**B**) % of EASs with protruding MVs on their surface. Three independent experiments in duplicate were performed and 100 parasites exhibiting at least one swelling were randomly counted per sample using SEM. Data are expressed as means ± SD. Approximately, 45% of parasites with flagellar swelling exhibited associated MVs.

### VPS32 localizes to the EASs and its overexpression increase EASs formation in *T. vaginalis* and *T. foetus*

ESCRTIII complex (Endosomal sorting complex required for transport) is a key player in the regulation of membrane fission during MVs formation and membrane remodeling (McCullough et al., 2018). VPS32 is an important component of the ESCRT-III complex (Cashikar et al., 2014). Hence, we transfected an HA-tagged version of the full length protein (VPS32FL-HA) in *T. vaginalis* and *T. foetus* to evaluate its localization by epifluorescence microscopy. Using an anti-HA antibody, we demonstrated that TvVPS32 and TfVPS32 are localized in the tip or along of recurrent and anterior flagella of parasites cultured in the absence of host cells (Fig. 10A). In concordance, the presence of TvVPS32 in the surface of extra-axonemal structures (EASs) as well as in MVs that protrudes from EASs was observed by immuno-gold electron microscopy using anti-HA antibody (Fig.10B). Based on this observation, we investigated the role of VPS32 EASs formation by analyzing the number of EASs in flagella of TvVPS32FL and TfVPA32FL parasites compared to parasites transfected with an empty plasmid (EpNeo). Interestingly, 15% and 18% of EASs were observed in TvVPS32 and TfVPS32 transfected parasites, respectively compared to 5% of EASs observed in EpNeo parasites (Figs.10C-D).

**Figure 10.**
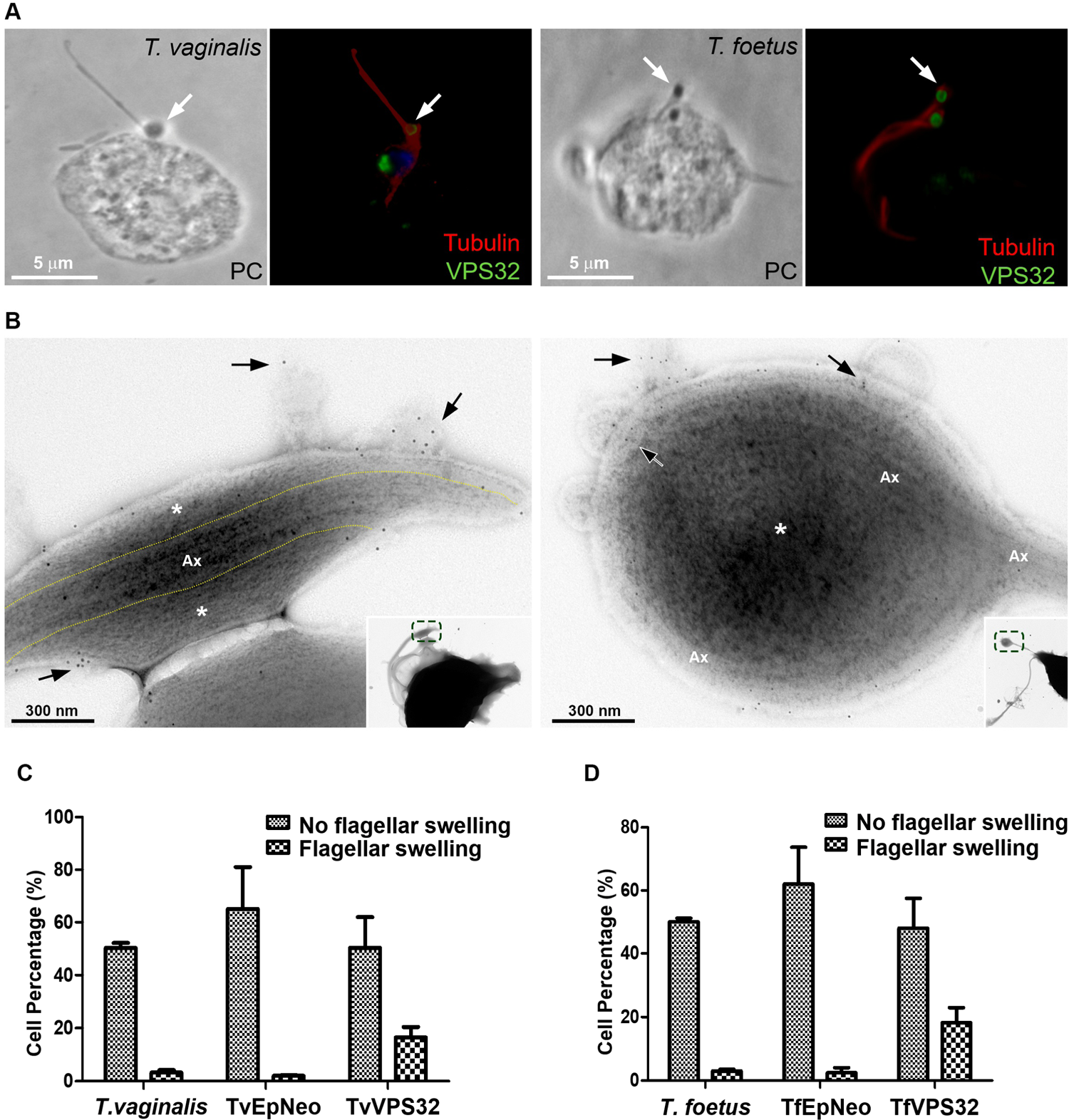
VPS32 is present in EASs membrane, and VPS32 overexpression increase EASs formation in *T. vaginalis* and *T. foetus*. (**A**) Representative immunofluorescence microscopy images. Cells exogenously expressing TvVPS32 and TfVPS32 with a C-terminal haemagglutinin (HA) tag were stained for immunofluorescence microscopy using a rabbit anti-HA antibody (green). PC, phase-contrast image. The nucleus (blue) was also stained with 4′,6′-diamidino-2-phenylindole (DAPI) and the flagella (red) was stained with mouse anti-tubulin antibody. Arrows indicate the flagellar subcellular localization of VPS32 in parasites cultured in the absence of host cells. (**B**) Negative staining of TvVPS32-HA transfected parasites immunogold-labelled with anti-HA antibodies demonstrate that TvVPS32 is localized in the surface of extra-axonemal structures (EASs) as well as in MVs that protrudes from EASs (arrows). (**C**) Analysis of the percentage of EASs in flagella of TvVPS32FL and TfVPA32FL parasites. Three independent experiments in duplicate were performed and 100 parasites exhibiting at least one swelling were randomly counted per sample using phase contrast microscope. Data are expressed as means ± SD. Approximately, 15% and 18% of flagellar EASs were observed in TvVPS32 and TfVPS32 transfected parasites, respectively compared to 5% of EASs observed in EpNeo parasites.

### TvVPS32 might regulate parasites motility

Information exchange between parasites of the same species could govern the decision to divide, to differentiate or to migrate as a group (Roditi, 2016). In some cases, this communication involves flagellar membrane fusion and the rapid exchange of proteins between connected cells (Szempruch et al., 2016). In this sense, our SEM observations demonstrate that *T. vaginalis* and *T. foetus* can connect to themselves by EASs present in flagella (Fig. 11A). Similarly, we observed that TvVPS32 transfected parasites can connect each other through the flagella and that TvVPS32 is localized in the flagella of parasites in contact (Fig. 11B). Based on this observation, we next decided to assess the motility capacity of TvVPS32 transfected parasites. To this end, TvEpNeo and TvVPS32 parasites were spotted onto soft agar and their migration capacity was analyzed by measuring the size of the halo diameter from the inoculation point to the periphery of the plate. As shown in Fig. 11C, the parasites transfected with TvVPS32 have a higher capacity of migration compared to parasites transfected with TvEpNeo, which might be suggesting a possible role for VPS32 protein in parasite motility.

**Figure 11.**
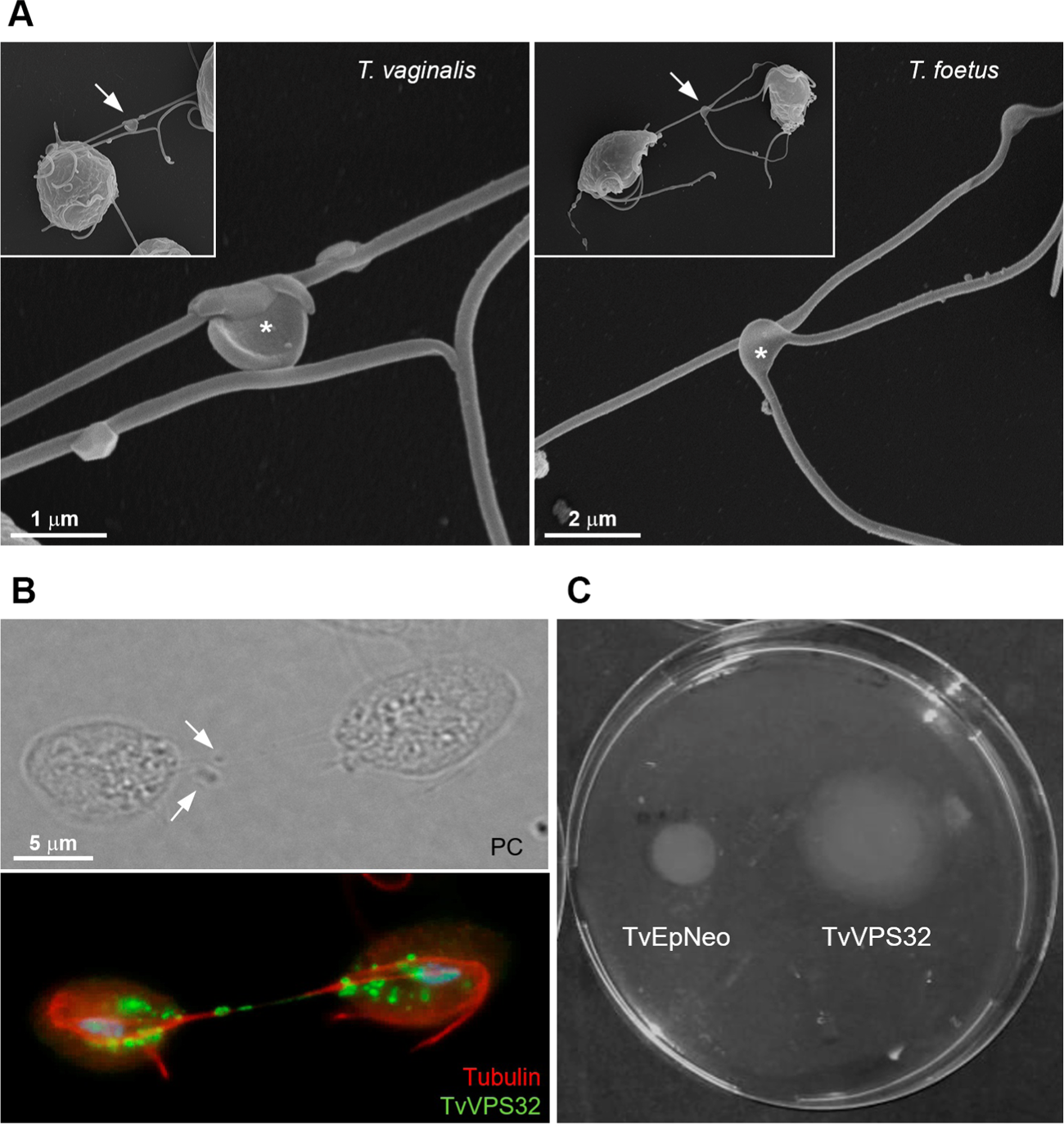
TvVPS32 plays a role in parasite motility. (**A**) Representative SEM images of parasites (*T. vaginalis* and *T. foetus*) connected to themselves by EASs (arrows). Notice the EASs in flagella that connect two parasites (*). (**B**) Immunofluorescence images showing that TvVPS32 transfected parasites connect each other through the flagella and that TvVPS32 is localized in the flagella of parasites in contact. TvVPS32 parasites cultured in the absence of host cells were co-stained with anti-HA (green) and tubulin (red). The nucleus (blue) was also stained with DAPI. Arrows indicate the EASs PC, phase-contrast image. (**C**) Representative TvVPS32 parasites motility assay. TvEpNeo and TvVPS32 parasites were spotted onto soft agar and their migration capacity was analyzed by measuring the size of the halo diameter during 4 days under microaerophilic conditions at 37°C. TvVPS32 parasites showed a higher capacity of migration compared to TvEpNeo parasites.

## DISCUSSION

Flagella have been extensively described as important players for host invasion, pathogenicity, and intercellular communication in pathogenic protists, mainly in kinetoplastids (Frolov et al., 2018; Kelly et al., 2020; Shimogawa et al., 2018). However, the structural organization and biological functions of trichomonads flagella remain largely unexplored. Most of studies about trichomonads flagella have focused on specializations of the flagellar membrane (Benchimol et al., 1982; Benchimol et al., 1992; Honigberg et al., 1984), propulsion force (Lenaghan et al., 2014; Ribeiro et al., 2000), and axoneme structure (Lee et al., 2009; Lopes et al., 2001; Melkonian, 1991). Here, we used a combination of electron microscopy techniques to reveal the ultrastructure of a novel extra-axonemal structure (EAS) in *T. vaginalis* and *T. foetus*, the most studied and important human and veterinary trichomonads, respectively. Traditionally, it has been assumed that *T. vaginalis* and *T. foetus do* not have EASs (Benchimol, 2004; Lenaghan et al., 2014; Melkonian, 1991; Rocha et al., 2010); however, we observed, in addition to the classical axoneme, thin fibrillary structures surrounded by the flagellar membrane running longitudinally along the axonemes. This novel structure displays morphology of paraflagellar swellings when seen by SEM or light microscopy. These EASs are more frequently found at the tip of the anterior and recurrent flagella in *T. vaginalis* and *T. foetus*, respectively. Suggesting that the EASs might be an evolutionarily conserved in the Parabasalia Phylum, the ultrastructural features of *T. vaginalis* and *T. foetus* EASs are similar to the extra-axonemal filaments described in other trichomonads and related parabasalid species, such as, *Trichomitus batrachorum* (Mattern et al., 1973), *Tritrichomonas muris* (Viscogliosi & Brugerolle, 1993), *Pentatrichomonoides* sp (Brugerolle & König, 1994), *Pseudotrypanosoma giganteum* (Brugerolle, 1999) and *Gigantomonas herculea* (Brugerolle, 2005).

Although the ultrastructure of *T. vaginalis* and *T. foetus* has been extensively investigated (Benchimol, 2004; de Andrade Rosa et al., 2013; de Souza & Attias, 2018), we believed there are some reasons that could explain why EASs had not been reported before. First, under axenic growth conditions, the EASs are only observed in 1-11% of parasites. Considering these percentages, a careful observation under electron microscope, mainly TEM, might be needed to be able to identify and properly investigate this structure. Second, as flagellar swellings can exhibit distinct morphologies, sizes, and relative positions, they may have been misinterpreted as a feature of cell death, i.e., flagellar blebbing, or an abnormality. Third, the EASs may have been considered as an artefact and just ignored or underappreciated by the investigators. In this regard, different authors using staining methods for light microscopy have described that the flagella of several parabasalids, including *T. vaginalis* and *T. foetus*, usually end with a granular or small swelling structure called “knob” (Čepička, 2016; Honigberg & King, 1964; Kirby, 1951); however, it has suggested that “knobs” may be artifacts due to the cell shrinkage during the fixation for protargol staining (Čepička, 2016; Ceza et al., 2015). Based on their location and morphologic similarities, we hypothesize that the EASs described here and the previously described “knobs” might be the same structure.

The EASs are found in the flagella of many cells including outer dense fibers and fibrous sheath of rodents and human sperm (Eddy et al., 2003; Linck et al., 2016), mastigonemes in *Chlamydomonas* (Liu et al., 2020), vane structures in the fornicate *Aduncisulcus paluster* (Yubuki et al., 2016), and the paraflagellar rod (PFR) of euglenoids and kinetoplastids (Zhang et al., 2021). They can run along the full length (outer dense fibers, fibrous sheath, and PFR), or just a portion, one or two-thirds of the axoneme (mastigonemes and vane structures). All those EASs have a striated appearance when viewed using TEM, suggesting a regular high-order structure. Similarly, the *T. vaginalis* and *T. foetus* EAS has also a striated fibrillar structure; however, whereas the outer dense fibers, mastigonemes, and PFR are regular intricate structures, linked to the axoneme via outer microtubule doublets, and found in all flagella from their respectively cell types (Linck et al., 2016; Liu et al., 2020; Zhang et al., 2021), the trichomonads EAS: (a) is not observed in all cells and axonemes; (b) it can be seen at the tip and/or middle of axoneme; (c) no association between the extra-axonemal filaments and axoneme microtubule doublets is still found; and (d) the organization and amount of the filaments can vary, resulting in two basic distinct morphologies, “sausage” and “spoon”, raging in different sizes. Those findings indicate that the assembly of trichomonads EAS is not a regular feature and might require cell signaling responses. Additionally, our results suggest that the several shapes and sizes of trichomonads EAS might correspond to different phases of a single assembly event. We hypothesize that the process might start with a “sausage” EAS and the “spoon” morphology might be the “final destination” morphology. Further analysis by videomicroscopy could help us to confirm this hypothesis. Moreover, we do not know yet whether the trichomonads flagellar swellings are reversible. Importantly, the identification of non-regular and transient EASs has never been described. Additional studies are needed to investigate the assembly kinetics and protein composition of trichomonads EAS.

The flagellum is a crucial host-pathogen interface, mediating attachment of parasites to host tissues (Kelly et al., 2020). In this regard, EASs, such as PFR, has an important role in flagellar retraction and flagellar support during tissues attachment in different stages of their life cycle (Bastin et al., 1996; Maga & LeBowitz, 1999; Maharana et al., 2015). Also, these PFR has been proposed as a metabolic, homeostatic, regulatory and sensory platform (Portman & Gull, 2010). These functions seem to be conserved among EASs during evolution. In this sense, the flagellar tip of *Crithidia fasciculata* is expanded up to six times its usual diameter upon contact with the insect host (Brooker, 1970). Also, arborescent outgrowths or “flagellipodia” were observed in the anterior flagellum of the bodonid flagellate *Cryptobia* sp. during their interaction to the snail *Triadopsis multilineata* (Current, 1980). Interestingly, the existence of flagellar morphological modifications seems to be related to adherence events along the life cycle in different flagellated organisms.

Here, we demonstrated that EASs formation increase during the attachment process in *T. vaginalis* and *T. foetus*. This finding is relevant considering that these protozoans are extracellular organisms, thus flagella and cell body are likely to play important roles in the initial adherence and survival of the pathogen on mucosal surfaces. It has been described that trichomonads flagella can interact with host epithelial cells, ECM proteins, yeasts, sperm cells and bacteria (Costa e Silva Filho et al., 1988; Midlej & Benchimol, 2010; Midlej et al., 2009; Pereira-Neves & Benchimol, 2007). *T. foetus* uses the recurrent flagellum to establish the first contact upon attachment with the host cell (Singh et al., 1999). Here, our results suggest a role of EASs during *T. vaginalis* and *T. foetus* attachment to the host cells as the EASs have been observed in direct contact with the host cells and the network-shaped mesh of preputial mucus. Taking into account that *T. vaginalis* appears to use its flagella as the guiding end to migrate and penetrate host tissues (Kusdian et al., 2013), we consider that structural changes due to EASs by increasing the adhesion surface would also facilitate trichomonads displacement in a viscous environment (epithelial mucus) or some materials (e.g. semi-solid media). However, future work is necessary to investigate this hypothesis.

In *T. vaginalis,* the EASs membranes possess high numbers of rosettes or intramembrane particles. The presence of intramembranous particles forming circular rosettes in the membrane of anterior flagellar of trichomonads has been previously reported (Benchimol & De Souza, 1990; Benchimol et al., 1981). The rosettes has been compared to particles involved in membrane fusion in *Tetrahymena* and hypothesized to contribute to active exo- and endocytosis (Lenaghan et al., 2014; Satir et al., 1973). These specialized integral membrane particles might be involved in active sensing of environment and play a key role in controlling local calcium levels to regulate flagellar beating (Benchimol, 2004; Lenaghan et al., 2014). In this sense, the kinetoplastid PFR provides a platform for cAMP and calcium signaling pathways that control motility, host–pathogen interactions, and for metabolic activities that may participate in energy transfer within the flagellum (Ginger et al., 2013; Portman & Gull, 2010; Shaw et al., 2019; Sugrue et al., 1988; Zhang et al., 2021). Similarly, the fibrous sheath of mammal sperm is a docking for key components in cAMP-signaling pathways, implicated in the regulation of sperm motility (Eddy et al., 2003). Based on the role of EASs in other organisms and our results, a sensory role for EASs might be suggested in *T. vaginalis*. The higher surface area of flagellar swellings due to EASs may provide a site for a greater number of rosettes.

The flagellar surface is a highly specialized subdomain of the plasma membrane, and flagellar membrane proteins are key players for all the biologically important roles of flagella (Landfear et al., 2015). In this sense, flagella are emerging as key players in cell-to-cell communication via shedding of microvesicles (MVs). MVs are observed protruding from flagellar tips of mammal cells (Nager et al., 2017; Salinas et al., 2017), the nematode *Caenorhabditis elegans* (Wang & Barr, 2018), and protists, including *Chlamydomonas* (Long et al., 2016), *Trypanosoma brucei* (Szempruch et al., 2016) and *T. vaginalis* (Nievas et al., 2018), suggesting that flagella may support MVs biogenesis. Here, we found MVs protruding from the trichomonads EASs. A higher area and curvature of the flagellar swellings may provide an advantage for the flagella to be used as a subcellular location for MVs biogenesis. In this context, ESCRT (endosomal sorting complexes required for transport) is an important mechanism known to facilitate the outward budding of membrane.

ESCRT proteins are emerging as versatile membrane scission machine that shape the behavior of membranes throughout the cell. In *Chlamydomonas reinhardtii*, ESCRT components are found in isolated ciliary transition zones, ciliary membranes, and ciliary microvesicles (Long et al., 2016). Additionally, ESCRT proteins mediate MVs release and influence flagellar shortening and mating (Diener et al., 2015; Long et al., 2016). ESCRT proteins are also found at the base of sensory cilia of *C. elegans* (Hu et al., 2007), suggesting that the ESCRT machinery are involved in flagellar function. In addition to mediate membrane budding and flagellar MVs shedding, ESCRT components may act as sensors for the generation and stabilization of membrane curvature of flagella (Jung et al., 2020; Long et al., 2016; Wang & Barr, 2018). Consistent with this, silencing of Vps36 in Trypanosomes, an ESCRT component, compromised the secretion of exosomes (Eliaz et al., 2017). In *T. vaginalis*, VPS32 protein (a member of ESCRT-III complex) has been identified in proteomic analyses of isolated exosomes and MVs (Nievas et al., 2018; Twu et al., 2013). Specifically, ESCRT-III has been shown to be crucial for diverse membrane remodeling events, the pinching off and release of MVs (Huber et al., 2020). Here, we revealed that VPS32 is present in the membrane as well as in MVs protruding from of EASs, in both *T. foetus* and *T. vaginalis*. Interestingly, we demonstrated that formation of paraflagellar swellings increase in parasites overexpressing TvVPS32 and TfVPS32; suggesting that ESCRT-III complex might be involved in EAS formation. Based on the function of ESCRT-III complex in other organism we could speculate that VPS32 might be regulating the dynamic flagellar membrane transformation that occurs during EASs formation. Alternatively, VPS32 could be participating in the final scission necessary for MVs release from flagellar membranes and subsequent membrane repair. Importantly, to our knowledge this is the first identification of an ESCRT protein associated with the flagella of a pathogenic protist.

In addition to release of extracellular vesicles, the contact between cells is also an important event in cell communication. Trypanosomes can interact with each other by flagellar membranes fusion, which could be partial and transient or irreversible and along the entire length of the flagellum (Imhof et al., 2016). These membrane fusion events might represent an alternative bidirectional mechanism used for proteins exchange with other individuals in a population. Fusion between membrane flagellar has been reported in *Crithidia Jasciculata* (Brooker, 1970)*, Leptomonas Iygaei* (Tieszen, 1989) and *Trypanosoma melophagium* (Molyneux, 1975). Curiously, in *Crithidia Jasciculata* has been described the existence of interflagellar type B desmosomes (temporary structures) between adjacent flagella of the microorganisms in contact to each other. Such junctions appear to maintain the “cluster” integrity that this protist form in the gut of the mosquito or in cultures (Brooker, 1970). The association of “clustering” and amoeboid transformation with a higher parasite adherence capacity has been reported in *T. vaginalis*, however, the mechanisms behind this phenomenon still remain unknown (Lustig et al., 2013). Here, we demonstrated that trichomonads can connect with each other by EAS flagellar, suggesting that this connection could contribute to cell communication. Supporting this, we observed that adhesion assays with Alcian blue- and fibronectin-coated coverslips induced amoeboid transformation, cell clusters (only Alcian blue), and increased the EAS formation, suggesting that could have a positive correlation between amoeboid transformation, cells clusters and EASs formation.

The results obtained here also demonstrated that TvVPS32 is present in EAS of parasites in contact to each other and interestingly, parasites overexpressing TvVPS32 showed greater motility in semisolid agar. Previously, we analyzed the growth rates of TvEpNeo and TvVPS32 parasites and we did not observe significant differences (data not shown); thus, an increase in halo size diameter is related to migration and not with increase parasites number. It has been reported that *Trypanosoma brucei* engages polarized migrations across the semisolid agarose surface mediated by flagellum communication (Oberholzer et al., 2010). Taking into account that VPS32 is the scission effector in different cellular membranes (Tang et al., 2015), we could speculate that this protein might be responsible of regulating different scission events during parasite: parasite communication or participating in flagellar membranes transformation important for parasite motility. However, future studies are needed to establish the specific function of ESCRT-III within this process in trichomonads.

This study will certainly shed light to our understanding on the flagella biology in pathogenic trichomonads. In summary, we described a novel EAS that provides a larger flagellar contact surface and added to this, the presence of rosettes and MVs in their membranes leads us to speculate that these structures could be involved in sensing, signaling, cell communication, and pathogenesis in trichomonads. In the future, continuing studies about the structure, proteomic, and assembly of EASs will enable us to better define how those mentioned functions are mediated by flagella in these extracellular parasites. Because the flagellum is an essential organelle, defining the flagellar morphology and roles in *T. vaginalis* and *T. foetus* may therefore help us to understand how the parasite colonizes the urogenital tract, how to prevent or treat infections, and uncover novel drug targets. In addition, trichomonads could emerge as a model system for studies of the conserved aspects of eukaryotic flagellum and EASs, providing new insights into evolutionary and functional aspects with direct relevance to other eukaryotes, including humans, in which flagella/cilia are essential for development and physiology, and defects can provoke several morbidities or fatal diseases.

## MATERIALS AND METHODS

### Parasites culture

The *T. vaginalis* strains B7RC2 (Parental, ATCC 50167), Jt and FMV1 (Midlej & Benchimol, 2010) and *T. foetus* K (parental) and CC09-1s strains (Pereira-Neves eta al., 2014) were cultured in Diamond’s Trypticase-yeast extract-maltose (TYM) medium supplemented with 10% bovine serum and 10 U/ml penicillin/10 ug/ml streptomycin (Invitrogen). Parasites were grown at 37°C and passaged daily. 100 μg ml−1 G418 (Invitrogen) was added to culture of the TvEpNeo/TvVPS32-HA and TfEpNeo/TfVPS32-HA transfectants.

### Plasmid construction and exogenous protein expression in trichomonads

The TvVPS32 construct was generated using primers with *NdeI* and *KpnI* restriction sites engineered into the 5′- and 3′-primers respectively. Polymerase chain reaction fragments were generated using standard procedures, and the resulting fragments were then cloned into the Master-Neo-(HA)_2_ plasmid to generate constructs to transfect into *T. vaginalis* and *T. foetus*. Electroporation of *T. vaginalis* G3 strain was carried out as described previously (Delgadillo et al., 1997), with 50 μg of circular plasmid DNA. Transfectants were selected with 100 mg ml−1 G418 (Sigma). The TfVPS32 construct was generated and transfected into *T. foetus* K as previously described (Iriarte et al., 2018).

### Scanning electron microscopy (SEM)

Cells were washed with PBS and fixed in 2.5% glutaraldehyde in 0.1 M cacodylate buffer, pH 7.2. The cells were then post-fixed for 15 min in 1% OsO4, dehydrated in ethanol and critical point dried with liquid CO2. The dried cells were coated with gold–palladium to a thickness of 25 nm and then observed with a Jeol JSM-5600 scanning electron microscope, operating at 15 kV.

### Transmission electron microscopy (TEM)

#### Routine preparation

The parasites were washed with PBS and fixed in 2.5% glutaraldehyde in 0.1 M cacodylate buffer, pH 7.2. The cells were then post-fixed for 30 min in 1% OsO_4_, dehydrated in acetone and embedded in Epon (Polybed 812). Ultra-thin sections were harvested on 300 mesh copper grids, stained with 5 % uranyl acetate and 1% lead citrate, and observed with a FEI Tecnai Spirit transmission electron microscope. The images were randomly acquired with a CCD camera system (MegaView G2, Olympus, Germany).

#### Negative staining

Parasites were settled onto positively charged Alcian blue-coated carbon film nickel grids (Labhart & Koller, 1981) for 30 min at 37°C. Next, cells were fixed in 2.5% glutaraldehyde in PEME (100 mM PIPES pH 6.9, 1 mM MgSO4, 2 mM EGTA, 0.1 mM EDTA) for 1h at room temperature. To better visualize the axoneme and EAS, parasites were permeabilized with 1% Triton X-100 for 10 min, washed with water, and negatively stained with 1% aurothioglucose (UPS Reference Standard) in water for 5 s. Alternatively, non-permeabilized cells were stained with 2% uranyl acetate in water for 10 s in order to visualize the flagellar rosettes. The grids were then air-dried and observed as described above.

#### Immunogold

Parasites were settled onto nickel grids as mentioned above, followed by fixation with 4% paraformaldehyde, 0.5 % glutaraldehyde in PEME for 1 h at room temperature. After washes in PEME, the grids were incubated with 1% Triton X-100 in PEME for 10 min and quenched in 50 mM ammonium chloride, 3% and 1% BSA, and 0.2% Tween-20 in PBS (pH 8.0). Next, the grids were incubated with anti-HA tag antibody (Invitrogen, 5B1D10), 10X diluted in 1% BSA in PBS for 3 h at room temperature. The grids were washed with 1% BSA in PBS and labelled for 60 min with 10 nm gold-labelled goat anti-mouse IgG (BB International, UK), 100 x diluted in 1% BSA in PBS, at room temperature. Samples were washed with PEME and water, negatively stained and observed as mentioned above. As negative control, the primary antibodies were omitted, and the samples were incubated with the gold-labelled goat anti-mouse antibody only. No labelling was observed under this condition.

### Parasite adhesion assays

Alcian blue and fibronectin were used in promoting cell adhesion to glass coverslips. Alcian blue-coated coverslips were prepared as previously described (Morone et al., 2006). Fibronectin-coated coverslips were prepared by first covering them with 100 µL of human (Sigma F0556) or bovine (Sigma F01141) fibronectin (working solution of 10 µg/mL in sterile PBS) for 1h at room temperature and washing them with sterile PBS. Parasites (1×10^6^cells/mL) were washed in PBS (pH 7.2) and resuspended in TYM medium without serum and PBS for Alcian blue and fibronectin assays respectively. A suspension of 50 µL was incubated on 1% Alcian blue or fibronectin-coated glass coverslips in humidity chamber for 0.5 to 2 h at 37°C. The parasites adhesion was monitored using an inverted phase contrast microscope. Non-adherent cells were collected with a pipette, harvested by centrifugation and washed with PBS. Next, the coverslips were rigorously washed with PBS to remove non-adherent parasites. Adherent cells remain on the coverslips even after several washes. Both adherent and non-adherent cells were then fixed and analysed using SEM as mentioned above. For the control experiments, parasites resuspended in TYM medium without serum or PBS were incubated on uncovered coverslips under the same conditions, collected with a pipette, harvested by centrifugation, and analysed as mentioned above.

### Parasite–host cell interaction

Human vaginal epithelial cells (VECs) were obtained from vaginal swabs of two healthy uninfected donors, 20 and 25 years old, with written consent. The cells were suspended in 20 ml of warm (37 °C) PBS (pH 7.2) just prior to experiments. Fresh bovine preputial epithelial cells (PECs) were kindly provided by Dr. Maria Aparecida da Gloria Faustino from the Faculty of Veterinary Medicine/Rural Federal University of Pernambuco. PECs were collected by aspiration with an artificial insemination pipette or by scraping the preputial cavity from a mature bull (> 4 years old) and suspended in 50 ml of warm (37 °C) PBS (pH 7.2) just prior to experiments. Next, VECs and PECs were washed 2 times in warm PBS by centrifugation at 400 x g for 5 min, suspended to a cellular density of 10^5^cells/ml in warm PBS and immediately used for interaction assays. VECs and PECs were co-incubated with *T. vaginalis* and *T. foetus*, respectively, at cell ratios of 1:1 or 5:1 parasite: host cell in PBS-F (PBS with 1% FBS at pH 6.5) at 37°C for 30 min. Prior to the co-incubation, parasites were washed three times in PBS, pH 7.2, and incubated to PBS-F at 37 °C for 15 min. In some assays, the human benign prostate epithelial line BPH1 was grown as described (Twu et al., 2013) and co-incubated with *T. vaginalis* as described above. For the control experiments, parasites incubated in PBS in the absence of host cells were analysed. The interactions were analysed using SEM, as mentioned above.

### Immunofluorescence assays

Parasites expressing the hemagglutinin-tag (HA) version of TvVPS32 and TfVPS32 were incubated at 37 °C on glass coverslips for 4 hours as previously described (Coceres et al., 2015). The parasites were then fixed and permeabilized in cold methanol for 10 min. Cells were then washed and blocked with 5% fetal bovine serum (FBS) in phosphate buffered saline (PBS) for 30 min, incubated with a 1:500 dilution of anti-HA primary antibody (Covance, Emeryville, CA, USA) and 1:500 dilution of anti-tubulin primary antibody diluted in PBS plus 2% FBS for 2 hours at RT, washed with PBS, and then incubated with a 1:5000 dilution of Alexa Fluor-conjugated secondary antibody (Molecular Probes) 1 hour at RT. The coverslips were mounted onto microscope slips using ProLong Gold antifade reagent with 4, 6′-diamidino-2-phenylindole (Invitrogen). All observations were performed on a Nikon E600 epifluorescence microscope. Adobe Photoshop (Adobe Systems) was used for image processing.

### Motility assay

Parasites TvEpNeo and TvVPS32 (1×10^6^ cells) were inoculated in soft-agar plates with Diamond’s, 5% FBS, and 0.32% agar and 10 U/ml penicillin/10 ug/ml streptomycin (Invitrogen). Parasites migration was monitored by analyze the colony diameter during 4 days under microaerophilic conditions at 37°C. Halo diameter was determined by ImageJ (image processing program).

### Quantitative analysis

The measurement of EAS filaments was carried out using TEM Imaging & Analysis (TIA) software of the microscope (FEI Company). The percentage of parasites that contain flagellar swelling was determined from counts of at least 500 parasites randomly selected per sample, using SEM or light microscope. The quantification of morphological aspects and distribution of flagellar swellings per cell was determined from counts of 100 parasites displaying at least one swelling per sample, using SEM. The morphology and relative position of flagellar swelling per flagellum was determined from counts of at least 100 anterior and recurrent flagella with swelling per sample, using SEM. The number of rosettes/µm^2^ was determined from counts of 50 flagella with or without swellings from at least ten randomly fields in the TEM grids using TIA software. The results are the average of three independent experiments performed at least in duplicate. Statistical comparison was performed ANOVA test, using computer analysis (GraphPad Prism v. 7.04, California, USA). *P<*0.05 was statistically significant.

## ACKNOWLEDGMENTS

We thank Dr. Marlene Benchimol from Universidade do Grande Rio for kindly providing *T. vaginalis* Jt and FMV1 strains and *T. foetus* K strain. We thank Dr. Milena Paiva and Dr. Maria Aparecida da Gloria Faustino from Instituto Aggeu Magalhães and Faculty of Veterinary Medicine/Rural Federal University of Pernambuco, respectively, for kindly providing PECs. We thank to Dr. Karina Saraiva and Dr. Cássia Docena from the Technological Platform Core of the Aggeu Magalhães Institute for their technical support.

## AUTHOR CONTRIBUTIONS

Conceived and designed the experiments: VMC, NdM, APN. Performed the experiments: VMC, LSI, AMM, TASA, APN. Analyzed the data: VMC, NdM, APN. Contributed reagents/materials/analysis tools: VMC, NdM, APN. Wrote the paper: VMC, NdM, APN. All the authors were involved in reviewing and editing the manuscript. All authors read and approved the final manuscript

---

**S1 Figure.**
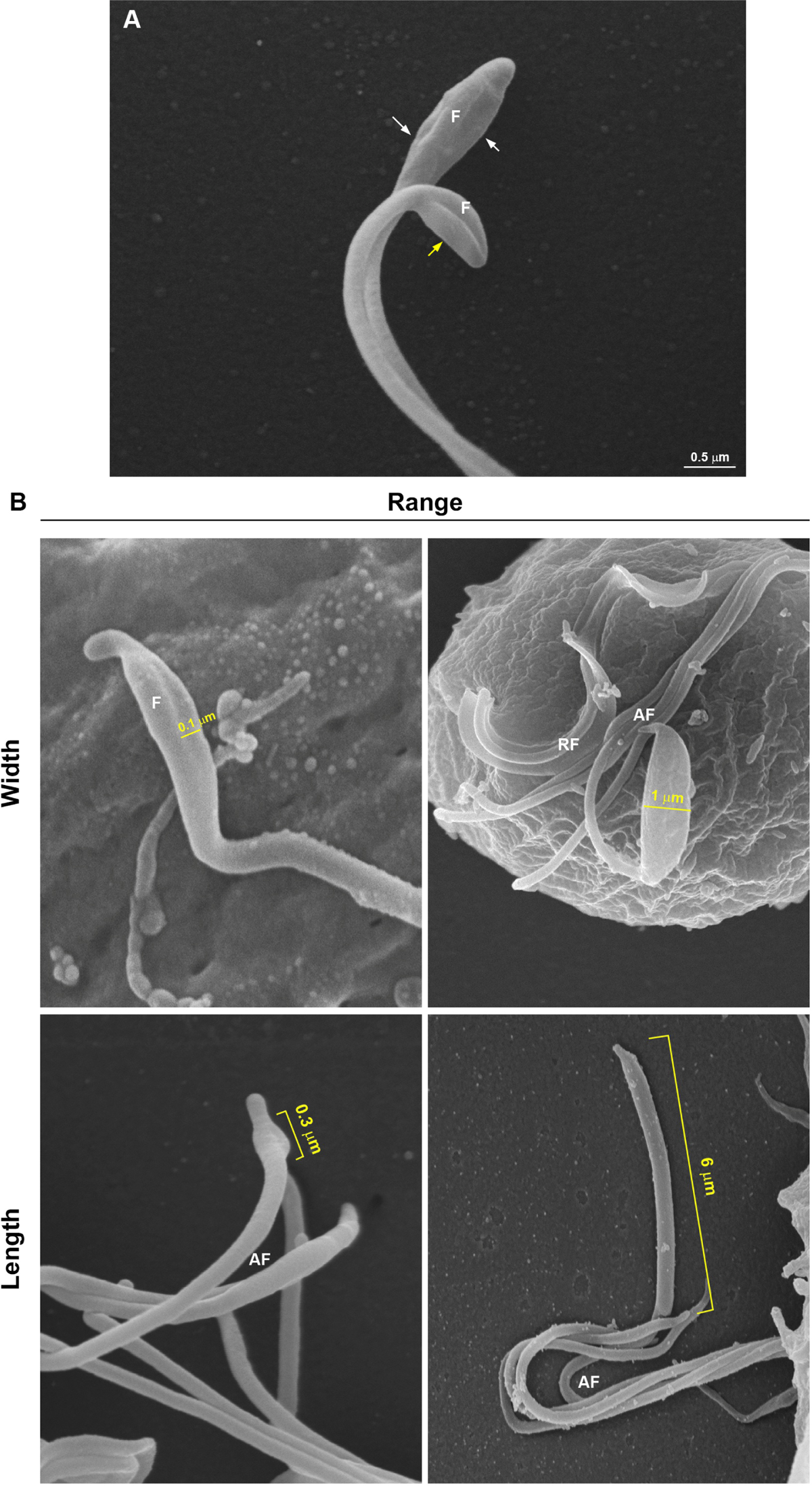
SEM of flagellar “sausage-like” swelling in *T. vaginalis*. (**A**) The swellings can be seen laterally (yellow arrow) to or surrounding (white arrows) the flagellum. (**B**) The swellings display a range size from 0.1 to 1 µm in thickness and a length from 0.3 to 6 µm. AF, anterior flagella; RF, recurrent flagellum.

**S2 Figure.**
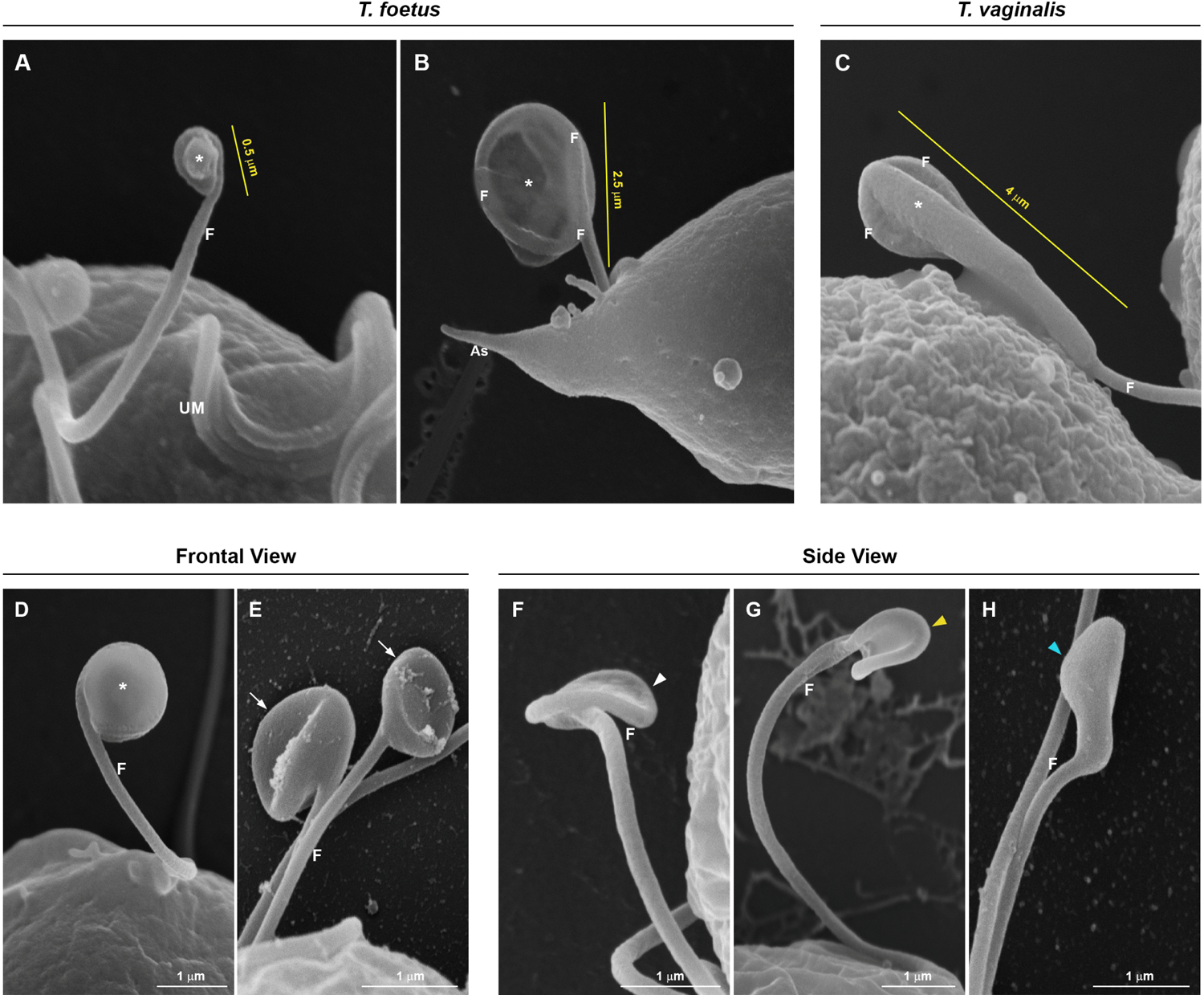
SEM of flagellar “spoon-like” swelling. The flagellum (F) folds around the swelling (*), forming a rounded or ellipsoid structure with a range size from 0.5 to 2.5 µm in the major axis in *T. foetus* (**A-B**) and more than 4 µm long in *T. vaginalis* (**C**). (**D-E**) Frontal views. The “spoon-like” structure exhibits a flattened (**D**) or concave (arrows) surface (**E**). (**F-H**) Side views. The structure (arrowheads) displays an aligned (**F**), curved (**G**), or convex (**H**) appearance.

**S3 Figure.**
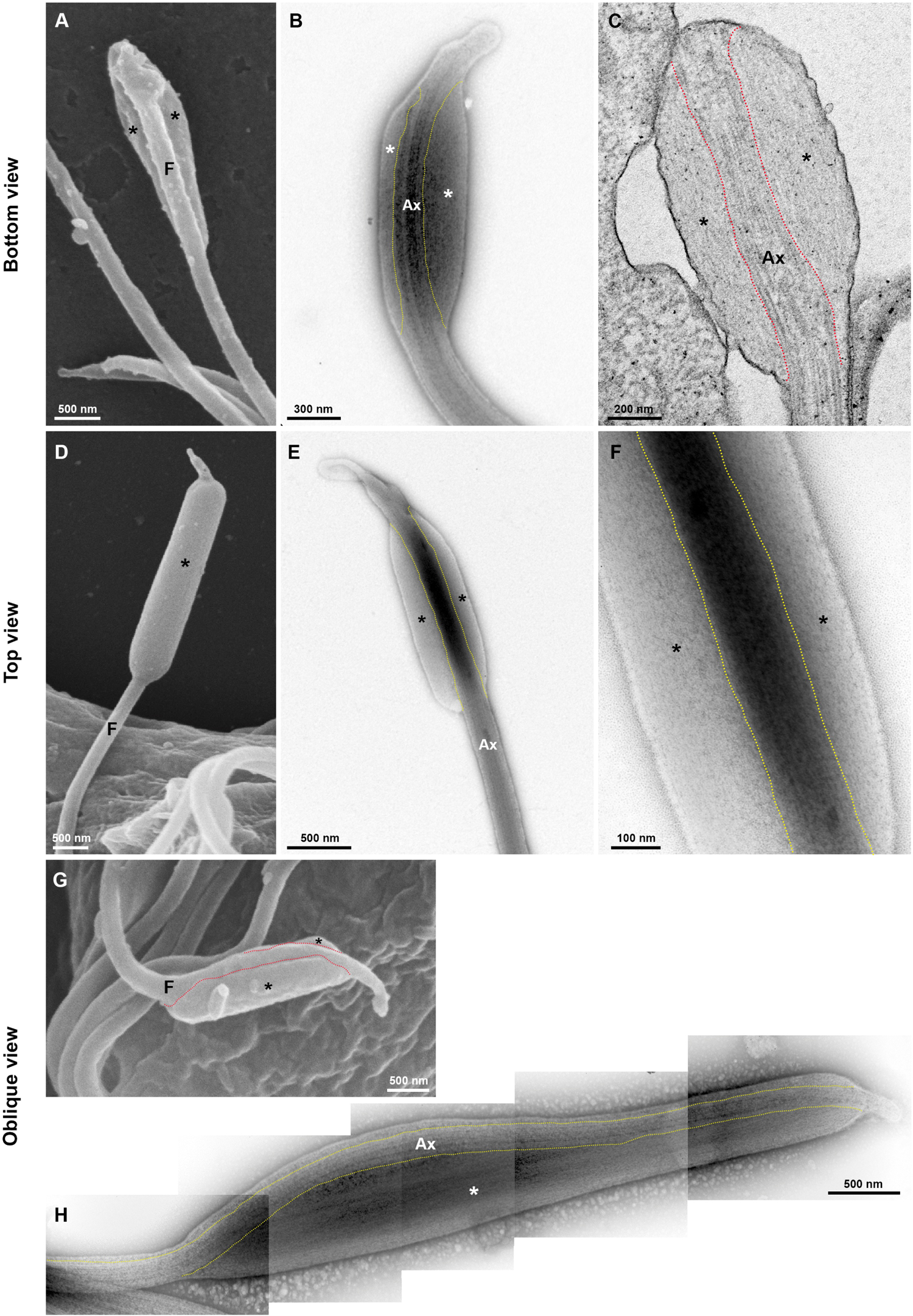
Ultrastructure of the sausage shaped swelling at the tip of *T. vaginalis* flagella seen by different perspectives. First row, SEM (**A**), negative staining (**B**) and ultrathin section (**C**) of swellings in a bottom view. Second row, SEM (**D**) and negative staining (**E-F**) of structure in a top view. Third row, SEM (**G**) and negative staining (**H**) of swelling in an oblique view. The dotted lines indicate boundary between axoneme (Ax) and the extra-axonemal filaments (*). In a SEM bottom view (**A**), notice that swelling (*) partially surround the flagellum (F), whereas in a top view (**D**) seems that the flagellum is totally surrounded by the swelling. In an oblique view (**G-H**), observe that the axoneme is in a slit of the swelling.

**S4 Figure.**
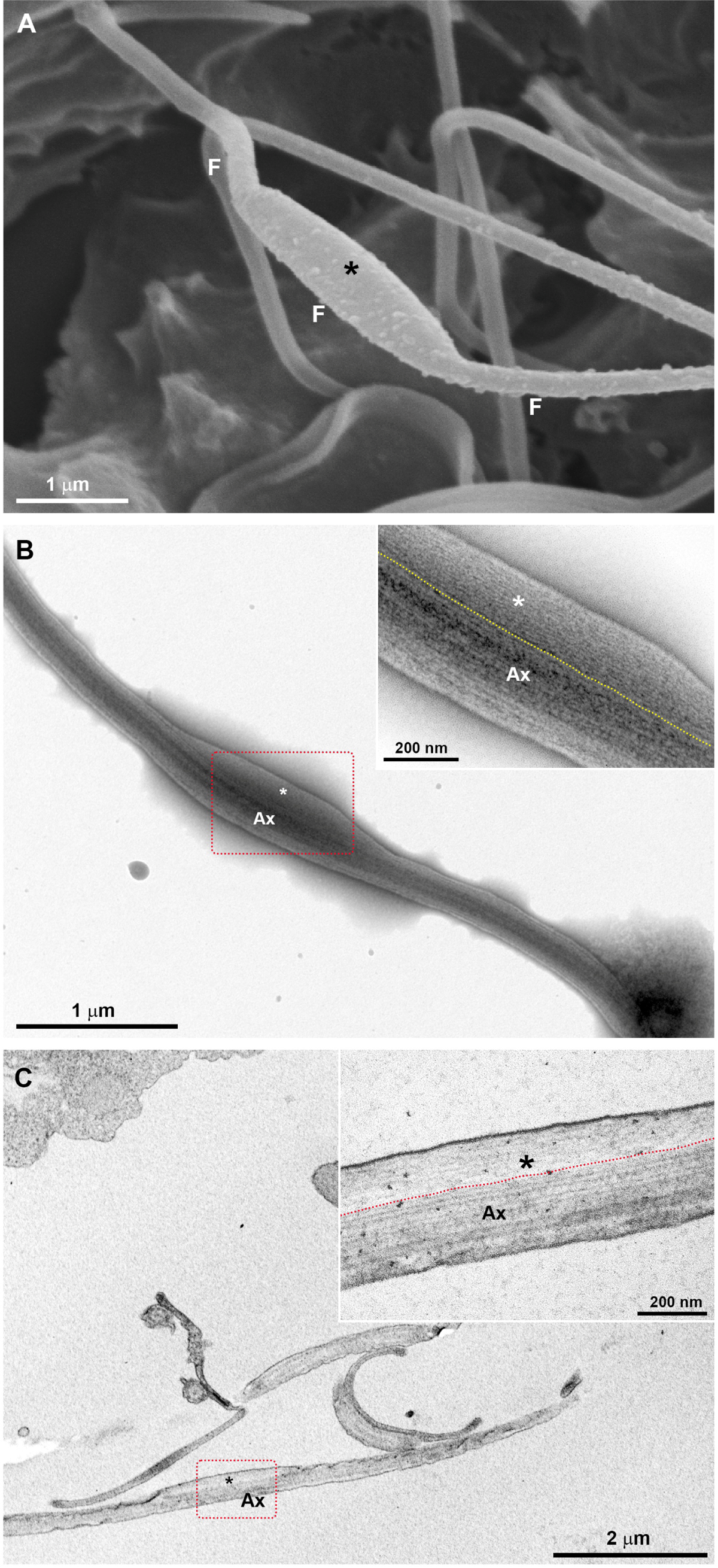
Fine structure of the “sausage-like” swelling in the middle of *T. vaginalis* flagella. (**A**) SEM. (**B**) Negative staining. (**C**) Ultrathin section. (**B-C**) The structure is formed by thin extra-axonemal filaments (*) that run longitudinally along the axoneme (Ax). The dotted lines indicate boundary between axoneme and the extra-axonemal filaments. F, flagellum.

**S5 Figure.**
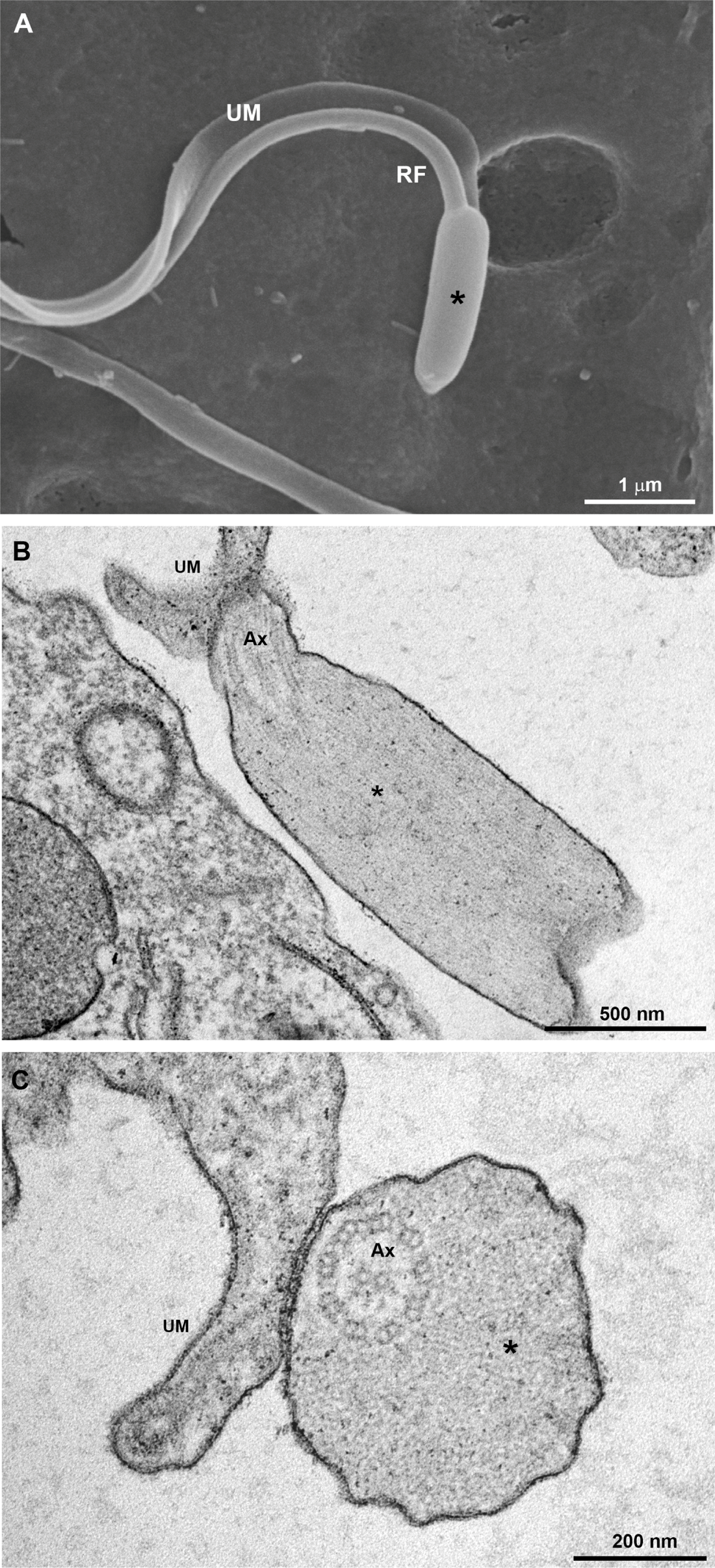
Ultrastructure of the sausage shaped swelling at the tip of *T. vaginalis* recurrent flagellum. (**A**) SEM. (**B**) Transversal and (**C**) cross sections. (**B-C**) The structure is formed by thin extra-axonemal filaments (*). RF, recurrent flagellum; UM, undulating membrane; Ax, axoneme.

**S6 Figure.**
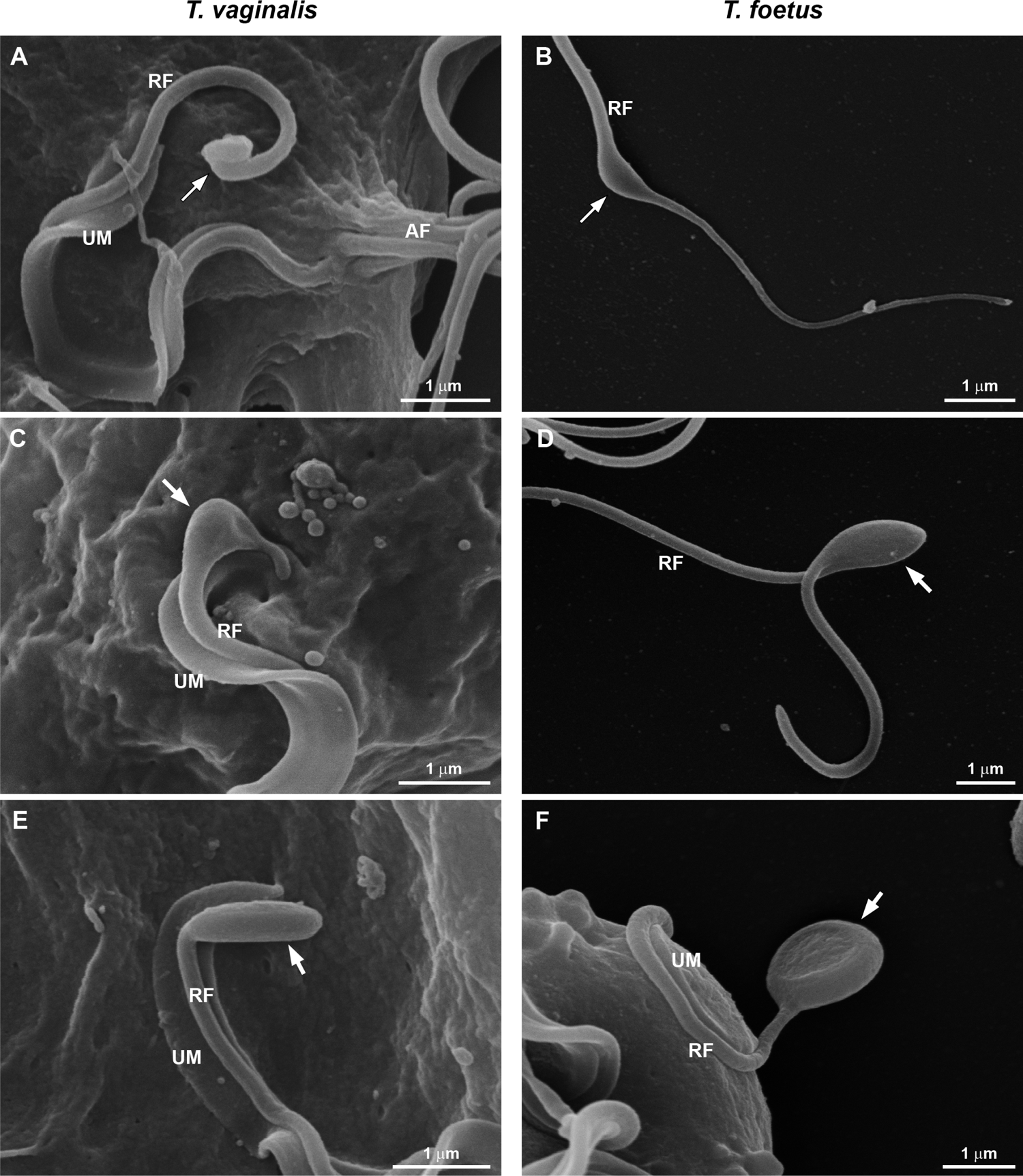
**SEM of flagellar swellings in the recurrent flagellum (RF) of *T. vaginalis* (A, C, E) and *T. foetus* (B, D, F)**. Notice that the *T. vaginalis*-RF has no free portion and exhibits sausage shaped swellings (arrows) of different sizes. Both sausage (**B**) and spoon (**D, F**) shaped swellings are seen in the free tip of *T. foetus*-RF. UM, undulating membrane.

**S7 Figure.**
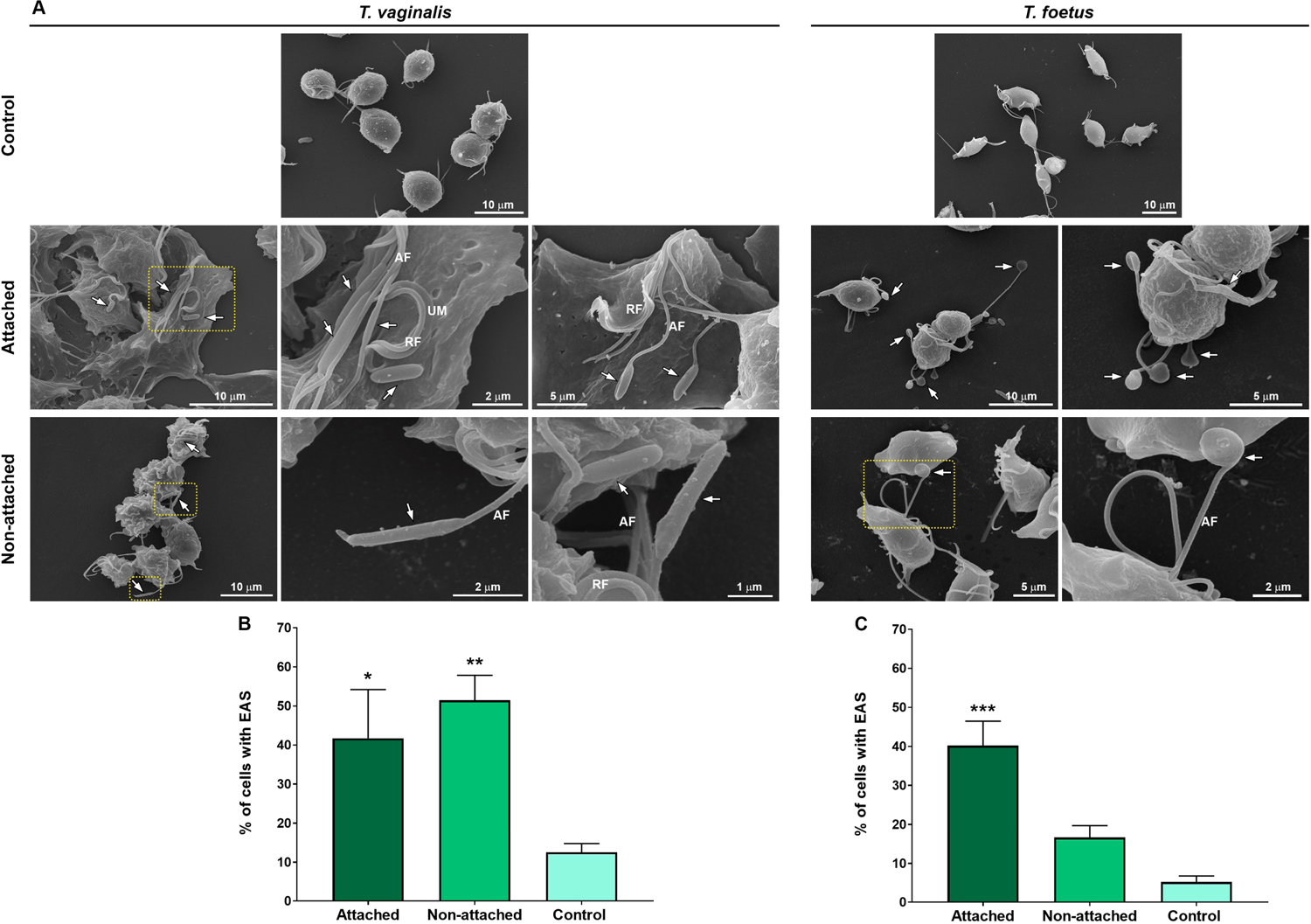
The EASs formation increase during trichomonads attachment on Alcian blue-coated coverslips. (**A**) Representative SEM micrographs of *T. vaginalis* and *T. foetus* after the adhesion assay. Control: parasites incubated on uncovered coverslips in humidity chamber for 0.5 h at 37°C, collected with a pipette, harvested by centrifugation, and prepared for SEM; Attached and non-attached: parasites incubated on 1% Alcian blue-coated glass coverslips in humidity chamber for 0.5 h at 37°C and rigorously washed with PBS to remove non-attached cells. Attached parasites remain on the coverslips even after several washes. Non-attached parasites were collected with a pipette, harvested by centrifugation, and prepared for SEM. “Control” is formed by non-adherent, suspended cells from uncovered coverslips, whereas non-adherent parasites from fibronectin are called “Non-attached”. In Control, the parasites display the typical pyriform cell body and no cell clusters. In Attached and Non-attached group, the cells are clustered, exhibiting an amoeboid or ellipsoid form and many flagellar swellings (arrows). AF, anterior flagella; RF, recurrent flagellum; UM, undulating membrane. (**B-C**) Quantitative analysis of the percentage of *T. vaginalis* (**B**) and *T. foetus* (**C**) with and without swelling after the adhesion assay. Three independent experiments in duplicate were performed and 500 parasites were randomly counted per sample using SEM. Data are expressed as percentage of parasites ± SD. The percentage of parasites displaying flagellar swelling in the Alcian blue-attached group is higher s when compared to control. Unexpectedly, the percentage of *T. vaginalis* with EAS in the Alcian blue-Non-attached group was significantly higher when compared to control. p<0.05; ** p<0.01; *** p<0.001 compared to control using One-Way ANOVA test (Kruskal-Wallis test; Dunn’s multiple comparisons test).

**S8 Figure.**
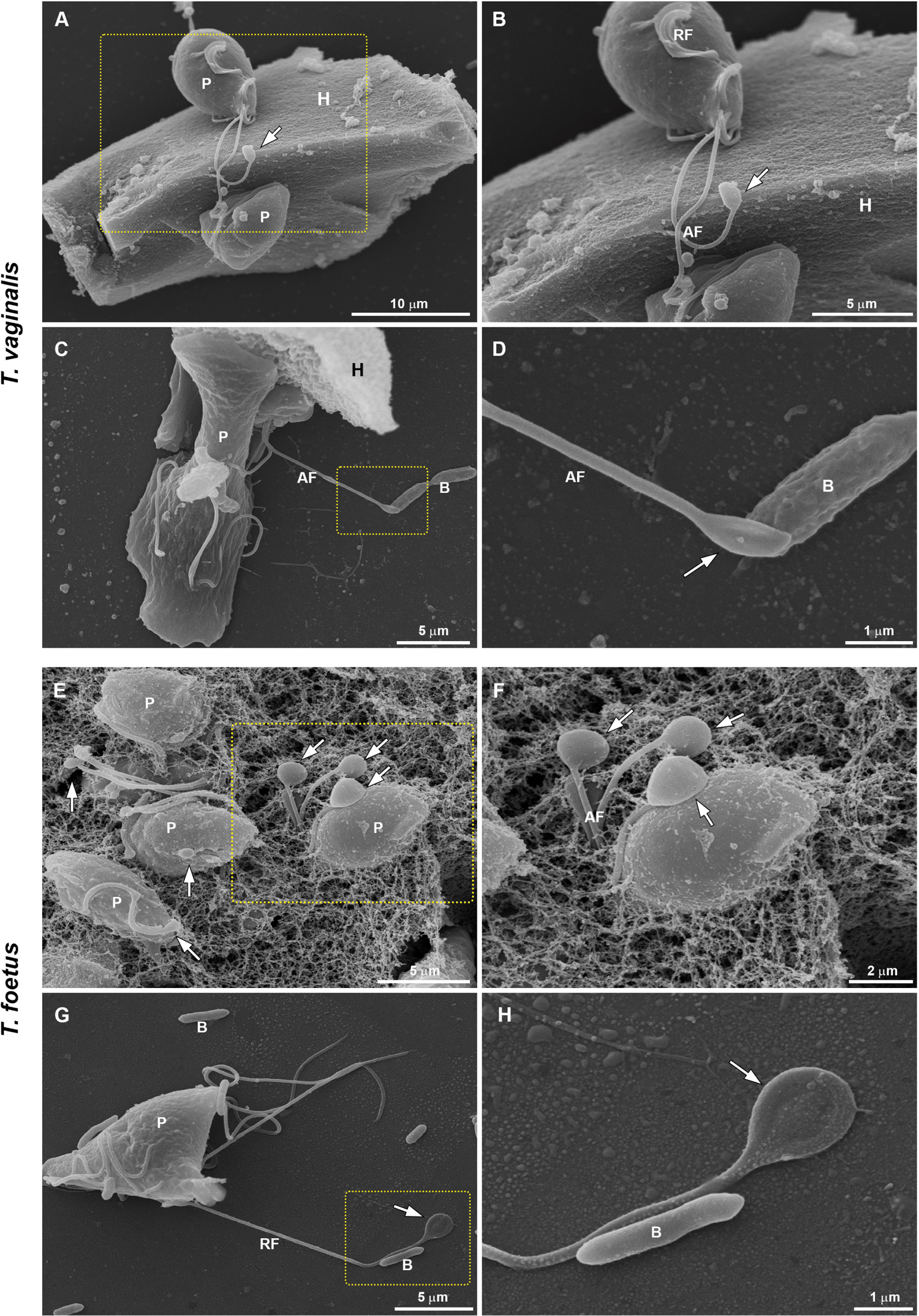
SEM of flagellar swellings in *T. vaginalis* and *T. foetus*. (**A-B**) Flagellar swelling (arrow) in a *T. vaginalis* (P) attached on a prostatic epithelial cell (H). (**C-D**) and (**G-H)** Swellings (arrows) in direct contact to bacteria (B) after host cells interaction assays. (E-F) Swellings (arrows) in *T. foetus* (P) adhered to the network-shaped mesh of preputial mucus. AF, anterior flagella; RF, recurrent flagellum.

Nome do arquivo:

https://d.docs.live.net/35fc6c14035a9c96/Documentos/Flagelos/Paper paraflagellar filaments/Figuras/Manuscript.docx Diretório:

Modelo:

C:\Users\anton\AppData\Roaming\Microsoft\Templates\

Normal.dotm

Título: Phagocytosis by Trichomonas vaginalis - New Insights Assunto:

Autor: User

Palavras-chave:

Comentários:

Data de criação: 26/07/2021 12:46:00

Número de alterações:107

Última gravação: 26/07/2021 23:17:00

Salvo por: Antonio Pereira

Tempo total de edição: 252 Minutos

Última impressão: 27/07/2021 00:22:00

Como a última impressão

Número de páginas: 54

Número de palavras: 13.339

Número de caracteres: 146.949 (aprox.)

